# A Computational Model of Tumor Interactions with Bone-Resident Cells Predicts Tumor-Type-Specific Responses to Perturbations

**DOI:** 10.64898/2026.02.16.706164

**Authors:** Alexandra Gutierrez Vega, Natalie E. Bennett, Erik P. Beadle, Saja Alshafeay, Ramcharan Chitturi, Anishka Nagarimadugu, Hrishikesh Villur, Arsh Jaiswal, Julie A. Rhoades, Leonard A. Harris

## Abstract

Tumor-induced bone disease (TIBD) arises from a complex interplay between metastatic cancer cells and the bone microenvironment, creating a self-reinforcing “vicious cycle” of bone destruction and tumor growth. Experimental evidence from our group (Buenrostro et al., *Bone* 113:77-88, 2018) suggests that tumor cells in the bone microenvironment early in disease rely more heavily on bone-derived growth factors, such as transforming growth factor-β (TGF-β), to sustain proliferation than tumor cells late in disease, which may grow independently of these factors. Here, we integrate a mechanistic, population-dynamics model of tumor–bone interactions with *in vivo* data to test the hypothesis that inhibiting bone resorption suppresses growth of non-adapted but not bone-adapted tumors. The model includes key regulators of TIBD, including TGF-β-driven tumor proliferation, parathyroid hormone-related protein (PTHrP) secretion, and osteoblast (OB)–osteoclast (OC) coupling. Parameter calibration using data from mice injected intratibially with parental (non-adapted) and bone-adapted breast cancer cells reveals distinct parameter values for each tumor type. Bone-adapted cells exhibit a higher basal division rate and reduced sensitivity to TGF-β-mediated stimulation, whereas parental-derived tumor cells depend more strongly on TGF-β and secrete PTHrP at higher rates to compensate for their slower growth. Model simulations reproduce the greater bone loss observed experimentally for bone-adapted tumors and predict that, for non-adapted tumors, bone destruction results from a slower but meaningful rise in OC activity and a possible moderate decline in OBs. Simulated treatment of bone-adapted tumors with the bisphosphonate zoledronic acid stabilizes bone density but has limited or highly variable effects on tumor growth. These results suggest that OC inhibition alone may be insufficient to restrain tumor expansion once tumors have adapted to the bone microenvironment. Together, these findings support the hypothesis that tumor adaptation to the bone microenvironment governs dependence on bone-derived growth factors and response to OC-targeted therapy, underscoring the value of mechanistic modeling for elucidating tumor–bone interactions and guiding tumor-type-specific treatment strategies for TIBD.

## Introduction

Tumor-induced bone disease (TIBD) arises from complex interactions between metastatic cancer cells and the bone microenvironment, creating a self-reinforcing loop of bone destruction and tumor growth, known as the “vicious cycle” (1–4). In healthy bone, homeostasis depends on tightly coupled interactions between osteoblasts (OBs), which form new bone, and osteoclasts (OCs), which resorb bone (5, 6). When tumor cells metastasize to bone, they disrupt this balance by secreting parathyroid hormone (PTH)-related protein (PTHrP) and other factors that stimulate OC activation and suppress OB function (4, 7). Activated OCs resorb bone, releasing growth factors, such as transforming growth factor-β (TGF-β), which, in turn, promote tumor proliferation and further PTHrP expression (2, 7). This feedback loop accelerates both tumor expansion and bone degradation, ultimately leading to osteolytic lesions, skeletal fragility, and significant patient morbidity (3).

Therapies that target OCs, such as bisphosphonates and receptor activator of nuclear factor-kappa (RANK) ligand (RANKL) inhibitors (8, 9), interrupt this cycle by suppressing bone resorption. These agents effectively preserve bone density and reduce skeletal-related events in patients, yet show limited impact on tumor burden and overall survival (8, 10). Experimental studies using *in vivo* models of bone metastasis (11, 12), such as intratibial tumor injection, genetically engineered mouse models, and patient-derived xenografts, have been reported that reflect these clinical patterns, i.e., bone-adapted tumors cause rapid bone loss that is only partially mitigated by OC inhibition (13, 14). For example, the initiation of OC inhibition early in the disease process, following intrafemoral tumor injection, has been demonstrated to be more efficacious than late-disease OC inhibition in reducing osteolytic lesions (15). However, data remains limited on the efficacy of OC inhibition for bone-adapted and non-adapted tumors. Addressing this question is crucial because if the efficacy of anti-resorptive treatments depends not only on their direct effects on OCs, but also on intrinsic tumor characteristics and differences in how tumors interact with and depend on the bone microenvironment, it could alter how TIBD is treated clinically.

In this study, we test the hypothesis that tumor cells not adapted to the bone microenvironment rely heavily on bone-derived growth factors, such as TGF-β, for proliferation, whereas bone-adapted cells can sustain growth through alternate means. To explore this, we integrate a mechanistic computational model of tumor–bone interactions with *in vivo* data from a murine model of bone-metastatic breast cancer using both intratibially injected bone-adapted and parental (non-adapted) tumor cells (11, 16). The computational model captures key regulatory processes in TIBD, including TGF-β-driven tumor proliferation, PTHrP secretion, and OB–OC coupling (5, 6, 17). Calibration of model parameters to experimental data reveals distinct differences between tumor types: tumors derived from bone-adapted cells exhibit higher basal division rates and reduced sensitivity to TGF-β-mediated stimulation, while tumors derived from parental cells depend more strongly on TGF-β and produce PTHrP at higher rates on a per-cell basis to compensate for their lower proliferation rate. Model simulations reproduce the greater bone loss observed experimentally for bone-adapted tumors and predict for non-adapted tumors that bone destruction results from a slower but meaningful rise in OCs and a possible moderate decline in OBs. Simulated treatment of bone-adapted tumors with a bisphosphonate stabilizes bone density but has limited or variable effects on tumor growth, suggesting OC inhibition alone may be insufficient to reduce proliferation once tumors adapt to the bone microenvironment.

Together, these findings provide a mechanistic explanation for tumor-type-specific differences in TIBD progression and therapeutic response. By quantitatively linking tumor adaptation to dependence on bone-derived growth factors, this work provides an explanation for why anti-resorptive therapies often fail for bone-adapted tumors despite effectively preserving bone (8, 10): non-adapted tumor cells rely heavily on bone-derived growth factors, such as TGF-β, to support proliferation, whereas bone-adapted tumors have intrinsic proliferative capacity and reduced dependence on TGF-β. As a result, inhibiting OCs and blocking the release of TGF-β interrupts the vicious cycle for non-adapted tumors but does not deprive bone-adapted tumor cells of essential proliferative signals. More broadly, this study illustrates the value of combining *in vivo* experimentation with computational modeling to disentangle complex cell–cell interactions in metastatic disease. The sections that follow describe the development of the mathematical model, calibration to experimental data, comparisons of parameter distributions between tumor types, and predictions of cellular dynamics and therapeutic outcomes under untreated and drug-treated conditions. We conclude with a discussion of the broader implications of this work for the study of TIBD.

## Results

### A Computational Model Integrating Tumor, OB, OC, and Bone Interactions

The computational model developed in this study integrates tumor growth, bone remodeling, and pharmacologic perturbation within a unified population-dynamics framework (Fig. 1). The core of the model is based on the bone homeostasis model introduced by Lemaire et al. (5), which describes the coupled dynamics of responding (i.e., inactive) OBs, active OBs, and active OCs regulated by PTH, RANK/RANKL/osteoprotegerin (OPG), and TGF-β signaling (Table S1, reactions 1–5). This regulatory module was reproduced exactly (*Materials and Methods*) and serves as the physiological baseline for all model extensions.

**Figure 1.**
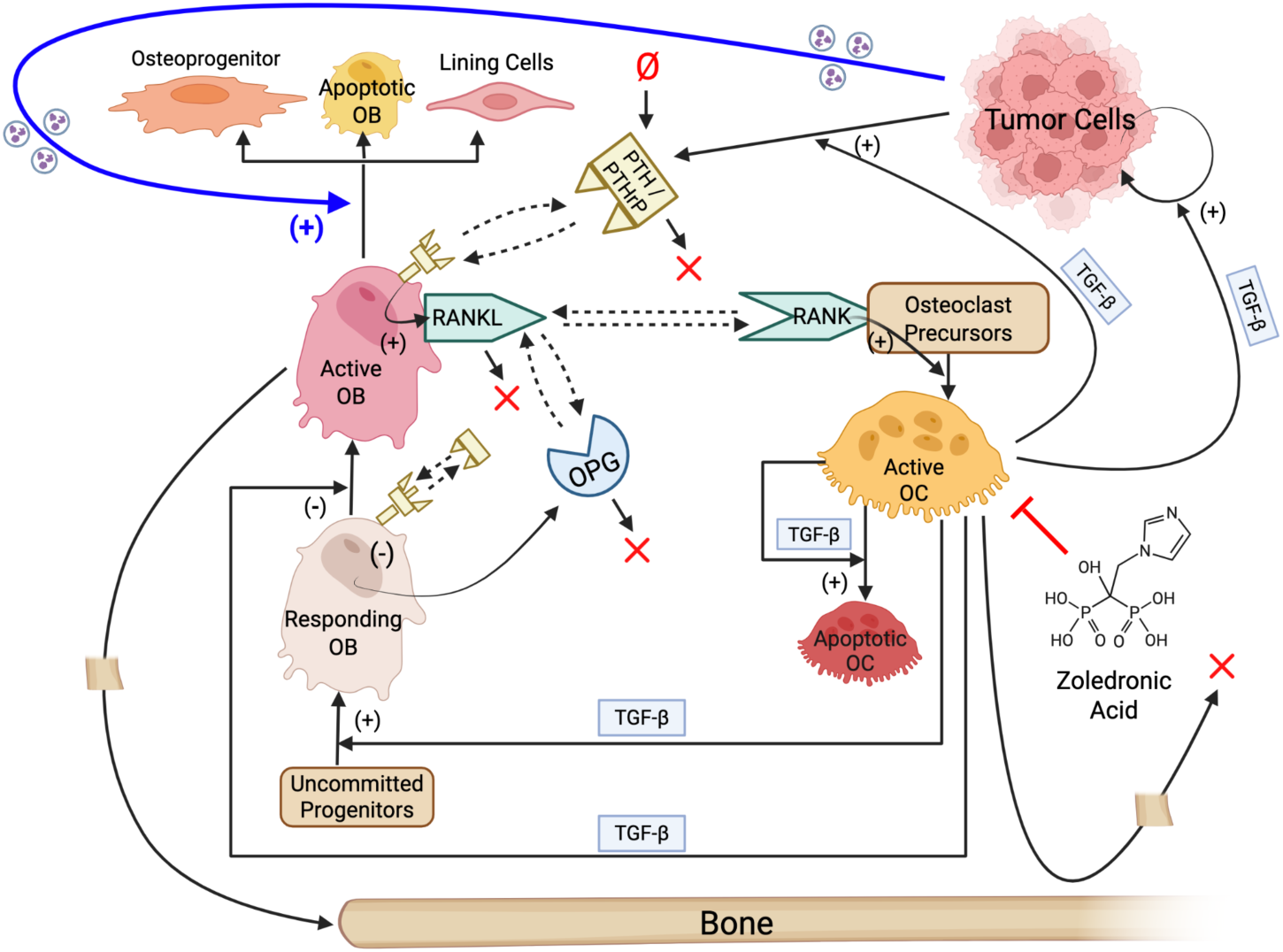
Model schematic illustrating interactions between tumor cells, osteoblasts, osteoclasts, and bone. The core of the model is taken from Lemaire et al. (5) and includes differentiation of, and interactions between, osteoblasts (OBs) and osteoclasts (OCs). Interactions involving tumor cells, bone, and the bisphosphonate zoledronic acid are added in this work. The thick blue arrow indicates a hypothesized effect of tumor-secreted factors on OB death (or differentiation to other cell types). (+): positive regulation; (-): negative regulation; ✕: degradation; Ø: synthesis; dashed arrows: binding/unbinding interactions; PTH: parathyroid hormone; PTHrP: PTH-related protein; OPG: osteoprotegerin; RANK: receptor activator of nuclear factor kappa B; RANKL: RANK ligand; TGF-β: transforming growth factor beta. Created with BioRender.com.

To adapt the Lemaire et al. (5) model to TIBD, bone density was introduced as an explicit dynamic variable with reactions describing bone synthesis by active OBs and bone resorption by active OCs (Table S1, reactions 6 and 7). Tumor cells were also incorporated, with reactions describing basal division, TGF-β–enhanced division, cell death, and density-dependent growth limitation via a carrying capacity (Table S1, reactions 8–10). Tumor-derived signaling was further incorporated through TGF-β–enhanced production of PTHrP by tumor cells (Table S2, function 4), as well as tumor-mediated active OB loss (death or differentiation) by an unspecified tumor-secreted factor (Table S1, reaction 11; blue arrow in Fig. 1). The latter assumption is supported by multiple studies reporting tumor suppression of OBs by secreted factors, such as activin A, sclerostin, tumor necrosis factor-α, and Fas ligand (18–20). Finally, pharmacological perturbation was incorporated through a reaction describing zoledronic acid (ZA)-induced death of active OCs (Table S1, reaction 12). This minimal representation reflects the primary, well-established mechanism of bisphosphonates as OC-targeted agents (8, 21, 22) and allows direct simulation of OC inhibition without assuming additional direct effects on tumor cells or OBs.

For bone formation and resorption (Table S1, reactions 6 and 7), Hill-type rate expressions (23, 24) were used that define effective *per-cell reaction rates* of the form

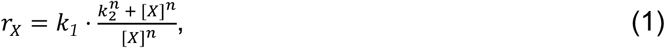

where *k_1_* and *k_2_* are constants, *n* is the Hill coefficient, and [*X*] denotes the population of active OBs or OCs (Fig. S1). The total reaction rate is obtained by multiplying Eq. (1) by the population(s) of the reactant species, i.e., bone formation scales with *r_B_*[*B*] and bone resorption with *r_C_*[*C*][*N*], where *B*, *C*, and *N* are active OBs, active OCs, and bone density, respectively. This formulation preserves biologically realistic limiting behavior: *r_X_* approaches zero as the cell population [*X*] vanishes, reduces to linear mass-action kinetics at high cell abundance (*r_X_* → *k*_1_ as [*X*] → ∞), and yields increased effective per-cell activity as OB and OC numbers decline (*r_X_* = 2*k*_1_ when [*X*] = *k*_2_). Thus, this model encodes an effective *homeostatic compensation* at the tissue scale in which OBs and OCs contribute more strongly to bone formation and resorption, respectively, when their population size is reduced. This is consistent with experimental and clinical observations that changes in OB and OC number do not scale linearly with net bone formation or resorption during pathological remodeling (2, 3, 10, 19, 21). Note that we initially attempted to model bone formation and resorption using standard mass-action kinetics. However, this formulation was unable to reproduce experimentally observed bone remodeling trajectories across tumor conditions.

Tumor cell division was modeled as the sum of a basal proliferation rate and a TGF-β–enhanced rate (Table S1, reaction 8), the latter of which depends on the quantity *π_C_* (Table S2, function 2). Inherited from the Lemaire et al. model (5), *π_C_* represents the fraction of TGF-β receptors on OB progenitor cells that are bound to TGF-β ligand. Practically, since TGF-β is not represented as an explicit species in the model, *π_C_* functions as a proxy for available TGF-β in the bone microenvironment, which regulates multiple processes, including responding OB production, OB activation, and OC death (Table S1, reactions 1, 2, and 5). In the present model, *π_C_* is further used to modulate tumor cell division (Table S1, reaction 8), encoding the assumption that elevated TGF-β signaling in the bone microenvironment enhances tumor proliferation (3, 25–27).

Tumor-derived PTHrP was incorporated into the model by modifying the PTH injection term, *I_P_*, from the Lemaire et al. model (5). PTH is not represented explicitly in the Lemaire et al. model (5) but influences bone remodeling through *I_P_* that appears in the ratio *π_P_* (Table S2, function 1), which, in turn, contributes to the function *π_L_* (Table S2, function 3) regulating OC production. To account for tumor-synthesized PTHrP, which we assume acts equivalently to PTH at the level of OB–OC regulation (28–30), we redefine *I_P_* as the sum of a constant baseline injection term and a tumor-dependent term that is proportional to the product of the tumor burden and the ratio *π_C_* (Table S2, function 4). This formulation is consistent with experimental evidence that TGF-β enhances PTHrP production and secretion by tumor cells (3, 31) and provides a mechanistic link between tumor burden, TGF-β signaling, and downstream bone remodeling feedback.

The final mechanistic model consists of six dynamical species, 12 reactions (Table S1), and four algebraic functions (Table S2). A total of 36 parameters appear in the model, of which 30 are treated as adjustable and estimated through calibration to experimental data (Table 1; *Materials and Methods*). This integrated computational framework provides a mechanistically grounded and quantitatively tractable platform for simulating tumor-induced bone remodeling and therapeutic intervention.

**Table 1.** Parameters for the population dynamics model of TIBD presented in this work. Parameters 1-23 are from the Lemaire et al. bone homeostasis model (5). Parameters 24-36 were added in this work. The six parameters annotated with an asterisk (*) in the first column were held fixed during parameter calibration. For additional context, see Tables S1 and S2 for the rate expressions and functions in which these parameters are included. *B*: active osteoblast (OB), *C*: active osteoclast (OC). Square brackets indicate concentration.

| Index | Parameter | Initial Value | Units | Description |
| --- | --- | --- | --- | --- |
| **Bone-Resident Cells** |  |  |  |  |
| 1 | $C^S$ | 0.005 | pM | Value of $[C]$ that gives $\pi_C = (1 + f_0)/2$ |
| 2 | $D_A$ | 0.7 | day<sup>-1</sup> | Rate of OC death enhanced by TGF-β |
| 3 | $d_B$ | 0.7 | day<sup>-1</sup> | Differentiation rate of responding OBs |
| 4 | $D_C$ | 0.0021 | pM·day<sup>-1</sup> | Differentiation rate of OC precursors |
| 5 | $D_R$ | 0.0007 | pM·day<sup>-1</sup> | Differentiation rate of OB progenitors |
| 6* | $f_0$ | 0.05 | unitless | Fixed proportion |
| 7* | $\bar{I}_L$ | 0 | pM·day<sup>-1</sup> | Constant rate of injected RANKL |
| 8* | $\bar{I}_O$ | 0 | pM·day<sup>-1</sup> | Constant rate of injected OPG |
| 9* | $\bar{I}_P$ | 0 | pM·day<sup>-1</sup> | Constant rate of injected PTH/PTHrP |
| 10 | $K$ | 10 | pM | Fixed concentration of RANK receptor |
| 11 | $k_1$ | 0.01 | pM<sup>-1</sup>·day<sup>-1</sup> | Rate of OPG-RANKL binding |
| 12 | $k_2$ | 10 | day<sup>-1</sup> | Rate of OPG-RANKL unbinding |
| 13 | $k_3$ | 0.00058 | pM<sup>-1</sup>·day<sup>-1</sup> | Rate of RANK-RANKL binding |
| 14 | $k_4$ | 0.017 | day<sup>-1</sup> | Rate of RANK-RANKL unbinding |
| 15 | $k_5$ | 0.02 | pM<sup>-1</sup>·day<sup>-1</sup> | Rate of PTH/PTHrP receptor binding |
| 16 | $k_6$ | 3 | day<sup>-1</sup> | Rate of PTH/PTHrP receptor unbinding |
| 17 | $k_B$ | 0.013 | day<sup>-1</sup> | Rate of active OB death |
| 18 | $K_L^P$ | $3 \times 10^6$ | unitless | Maximum number of RANKL attached on each cell surface |
| 19 | $k_O$ | 0.35 | day<sup>-1</sup> | Rate of elimination of OPG |
| 20 | $K_O^P$ | $2 \times 10^5$ | day<sup>-1</sup> | Minimal rate of production of OPG per cell |
| 21 | $k_P$ | 86 | day<sup>-1</sup> | Rate of PTH/PTHrP degradation |
| 22 | $r_L$ | 1000 | pM·day<sup>-1</sup> | Net rate of RANKL production and degradation |
| 23 | $S_P$ | 250 | pM·day<sup>-1</sup> | Rate of synthesis of systemic PTH |
| **Bone** |  |  |  |  |
| 24 | $k_{1B}$ | $3.36 \times 10^6$ | day<sup>-1</sup> | Minimum rate of OB production of bone |
| 25 | $k_{2B}$ | 0.1 | pM | Value of $[B]$ at which the OB bone production rate = $2k_{1B}$ |
| 26* | $n_B$ | 1 | unitless | Hill coefficient for OB bone production |
| 27 | $k_{1C}$ | $10^6$ | day<sup>-1</sup> | Minimum rate of OC consumption of bone |
| 28 | $k_{2C}$ | 0.001 | pM | Value of $[C]$ at which the OC bone consumption rate = $2k_{1C}$ |
| 29* | $n_C$ | 1 | unitless | Hill coefficient for OC bone consumption |
| **Tumor Cells** |  |  |  |  |
| 30 | $k_{Tdiv}$ | 1 | day<sup>-1</sup> | Basal rate of tumor cell division |
| 31 | $k_{Tgfb}$ | 0.1 | day<sup>-1</sup> | Rate of tumor cell division enhanced by TGF-β |
| 32 | $k_{Tdeath}$ | 0.6 | day<sup>-1</sup> | Rate of tumor cell death |
| 33 | $M$ | 0.9 | pM | Tumor cell carrying capacity |
| 34 | $k_{Tpthrp}$ | 500 | day<sup>-1</sup> | Rate of PTHrP production by tumor cells |
| 35 | $k_{T-B}$ | 2 | pM<sup>-1</sup>·day<sup>-1</sup> | Rate of OB death/differentiation enhanced by tumor cells |
| **Drug Treatment** |  |  |  |  |
| 36 | $k_{Z-C}$ | 1 | day<sup>-1</sup> | Rate of OC death by zoledronic acid |

### Establishment of a Murine Intratibial Model of Bone Metastasis

To quantify tumor-driven bone remodeling and generate experimental data for computational model calibration, we employed a previously published intratibial injection model of advanced bone metastatic disease (11, 12) (Fig. 2A). Bone-metastatic clones of MDA-MB-231 breast cancer cells were injected directly into the tibial marrow cavity, and disease progression was monitored longitudinally over 21 days (*Materials and Methods*). This model enables controlled initiation of tumor growth within the bone microenvironment and recapitulates key features of TIBD, including osteolysis, altered bone-resident cell populations, and progressive tumor expansion. Radiographic imaging was used to assess the emergence and progression of osteolytic lesions over time (Fig. S2), providing an initial readout of tumor-induced bone damage and informing downstream inclusion criteria for quantitative analyses. Micro–computed tomography (µCT) imaging reveals time-dependent bone destruction following tumor injection, with representative µCT reconstructions showing progressive loss of trabecular bone. Early surface irregularities are evident by Day 7 and extensive structural degradation by Day 21 (Fig. 2B). These imaging results confirm rapid and aggressive bone remodeling in the presence of bone-metastatic tumor cells and provide a quantitative measure of tumor-induced bone loss.

**Figure 2.**
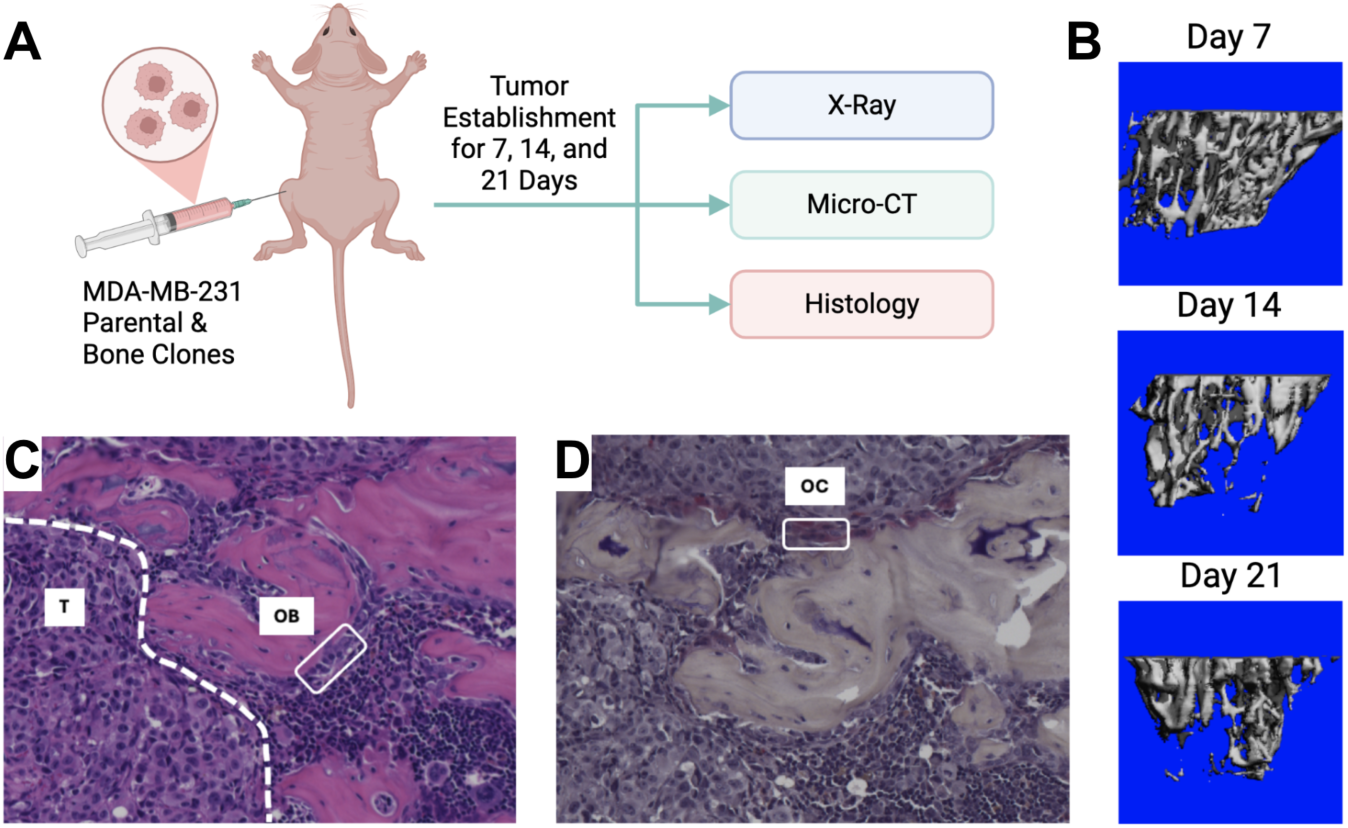
Murine model of bone metastasis establishment. **(A)** Female nude mice were injected intratibially with 50,000 MDA-MB-231b cells (bone-adapted), and disease was followed for 7, 14, and 21 days after injection. At each timepoint, hindlimbs were x-rayed and then collected for micro-computed tomography (µCT) and histology. **(B)** Representative 3D reconstruction of bone segments analyzed by µCT. **(C)** Representative H&E image showing a row of pale osteoblasts (OB) along the bone surface and established tumor (T) in the marrow space. **(D)** Representative TRAP stain slide showing a multinucleated osteoclast (OC) on the bone surface. H&E: hematoxylin and eosin; TRAP: tartrate-resistant acid phosphatase. Panel *(A)* created with BioRender.com.

Histological analysis further characterizes the cellular basis of this remodeling. Hematoxylin and eosin (H&E) staining demonstrate extensive tumor infiltration within the marrow space and marked disruption of normal bone architecture (Fig. 2C). OBs lining mineralized bone surfaces are substantially reduced in metastatic lesions, consistent with suppression of bone formation. In contrast, tartrate-resistant acid phosphatase (TRAP) staining reveals abundant multinucleated OCs localized to resorbing bone surfaces (Fig. 2D), indicating elevated OC activity as a primary driver of bone degradation in this model. Together, the imaging and histological data generated in this work establish a robust and biologically relevant *in vivo* platform for quantifying tumor-induced bone destruction and associated changes in bone-resident cell populations. These data form the foundation for subsequent model calibration and analysis and enable direct investigation of the mechanisms underlying tumor-type-specific bone remodeling dynamics.

### Monte Carlo-Based Model Calibration Accurately Captures In Vivo Tumor, Bone, and Bone-Resident Cell Dynamics

*In vivo* experiments performed in this study use bone-adapted MDA-MB-231 breast cancer cells (*Materials and Methods*) to quantify tumor-driven bone remodeling under untreated conditions (Fig. 3, *blue circles*). These data illustrate coordinated temporal changes in bone density (Fig. 3, *top left*), OC and OB abundances (Fig. 3, *top right* and *bottom left*, respectively), and tumor burden (Fig. 3, *bottom right*). Of note, bone density declined rapidly over the 21 days of the experiment, reaching approximately half its initial value by the end, and was coincident with a pronounced expansion of OCs and depletion of OBs. Tumor burden increased sharply over the same time frame, reflecting the aggressive growth behavior of bone-adapted breast cancer variants. Together, these measurements provide a comprehensive view of metastatic tumor–bone interactions *in vivo* and serve as the primary constraint for calibrating tumor–bone coupling dynamics in the computational model.

**Figure 3.**
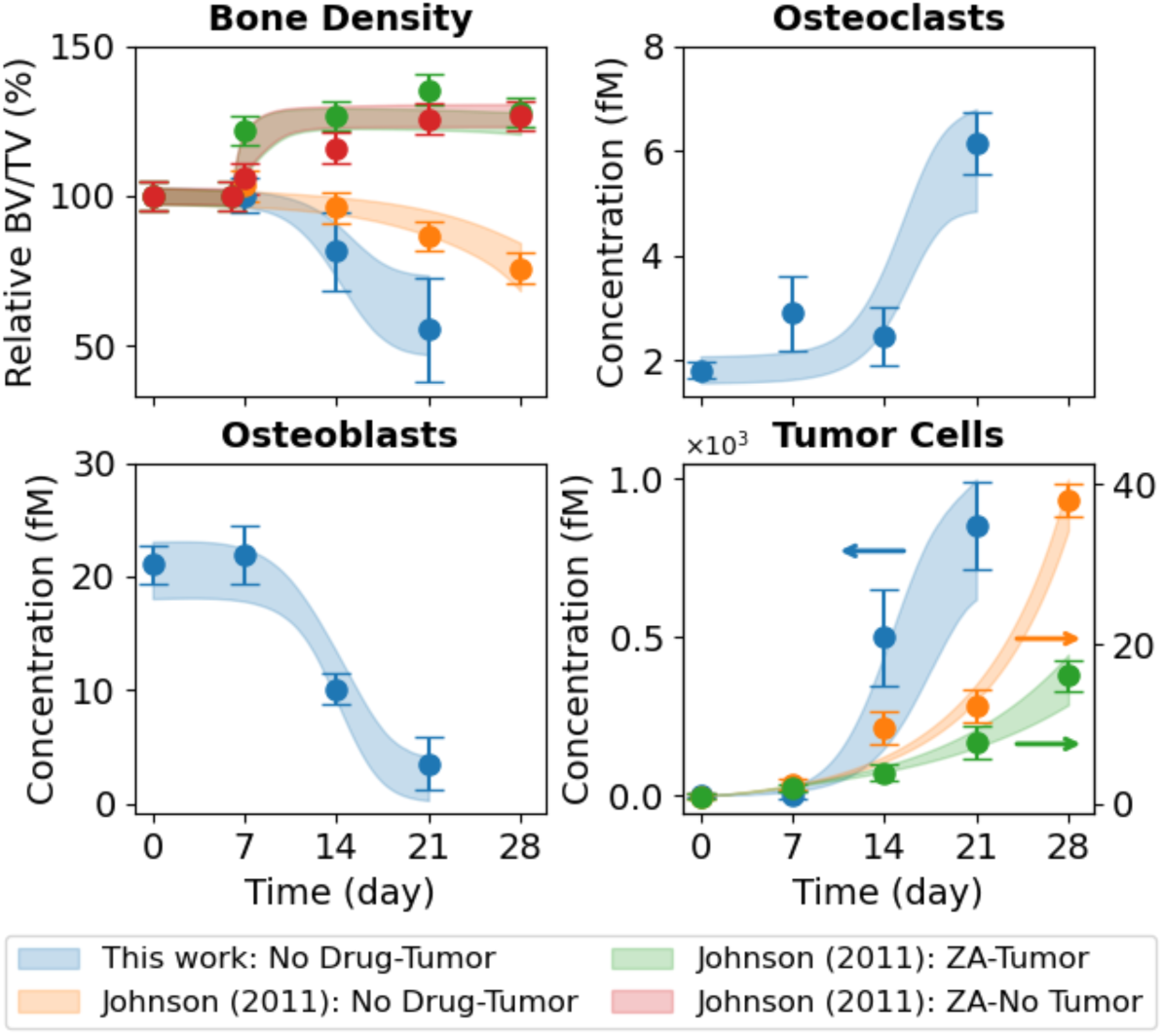
Monte Carlo-based model fits to *in vivo* experimental data. The mathematical model was fit to nine sets of experimental data simultaneously. For the bone-adapted tumor without drug treatment (“This work: No Drug-Tumor”), experimental measurements are shown for bone density, osteoclasts (OCs), osteoblasts (OBs), and tumor cells. For the parental-derived tumor, bone density and tumor data are from Johnson et al. (16) for three experiments: tumor without drug treatment (“Johnson (2011): No Drug-Tumor”), tumor with drug treatment (“Johnson (2011): ZA–Tumor”), and drug treatment without tumor (“Johnson (2011): ZA-No Tumor”). Experimental error bars are standard errors (σ/√n) and model fits are shown as shaded 90% confidence envelopes. Bone density is plotted as a percentage relative to Day 0. BV/TV: bone volume ÷ tumor volume; ZA: zoledronic acid.

In addition to the bone-adapted data generated in this study, we incorporated datasets reported by Johnson et al. (16) directly into the analysis (Fig. 3, *top left* and *bottom right*). These data were generated using parental (i.e., non-bone-adapted) MDA-MB-231 cells, which are distinct from the bone-adapted MDA-MB-231 variant used in the present work. The data points, which include bone density and tumor burden measurements for untreated tumor-bearing mice and tumor-bearing mice treated with the bisphosphonate ZA, and bone density measurements for ZA-treated mice without tumor, were extracted directly from the original publication (16). To facilitate model calibration, fluorescence tumor data were converted to estimated cell counts (Fig. S3), and error bars for both the tumor and bone density data were strategically modified (Fig. S4). Importantly, these modifications preserve the reported means while facilitating parameter identifiability (see *Materials and Methods* for details). The primary motivation for incorporating these datasets in this study was to enable estimation of the parameter value for ZA-mediated OC inhibition in the computational model (*k_Z–C_*; Tables 1 and S1). This interaction is assumed to be independent of tumor type and not identifiable from untreated metastatic tumor data alone.

Parameter estimation was performed using PyDREAM (32) (*Materials and Methods*), which inferred an ensemble of parameter sets that produce simulation outputs consistent with the bone-adapted data generated here and the three non-adapted tumor datasets from Johnson et al. (16) (Fig. 3). In all, the computational model (Fig. 1) was simultaneously fit to nine *in vivo* datasets from four independent experiments involving two tumor types injected intratibially with either bone-adapted or parental MDA-MB-231 breast cancer cells. Convergence of the parameter calibration process is evident by the stabilization of the inferred log-likelihood values for all Monte Carlo chains (Fig. S5) and quantified via the Gelman-Rubin metric (32, 33) (*Materials and Methods*). Posterior parameter distributions illustrate uncertainties in the parameter values, which vary significantly across parameters (Fig. S6). For all conditions, 90% confidence envelopes are consistent with the magnitudes of the experimental error bars (Fig. 3), indicating the inferred parameter ensemble captures observed biological variability/uncertainty.

### Tumor-Type-Specific Parameter Distributions Differentiate Non-Adapted and Bone-Adapted Tumor Dynamics

Since the experiments in Johnson et al. (16) use non-adapted cells from the parental MDA-MB-231 breast cancer cell line rather than from the bone-adapted version used in this study, to correctly perform parameter calibration we had to independently fit tumor-associated model parameters to the different datasets based on tumor type. Specifically, five parameters (*k_Tdiv_*, *k_Ttgfb_*, *k_Tdth_*, *k_Tpthrp_*, *k_T–B_*; see Table 1 and Fig. S6A,B) were fit independently to the bone-adapted tumor data from this work and the parental-derived tumor data from Johnson et al. (16). The other 25 shared parameters (Fig. S6C), including the ZA–OC interaction rate constant (*k_Z–C_*), were inferred jointly across all datasets. Values for these five parameters for the two different tumor types were compared by overlaying the inferred histograms from the PyDREAM calibration (Fig. 4A). Differences between histograms were quantified statistically using the histogram distance (34, 35) (Fig. 4B; *Materials and Methods*), which quantifies the degree of separation between parameter distributions.

**Figure 4.**
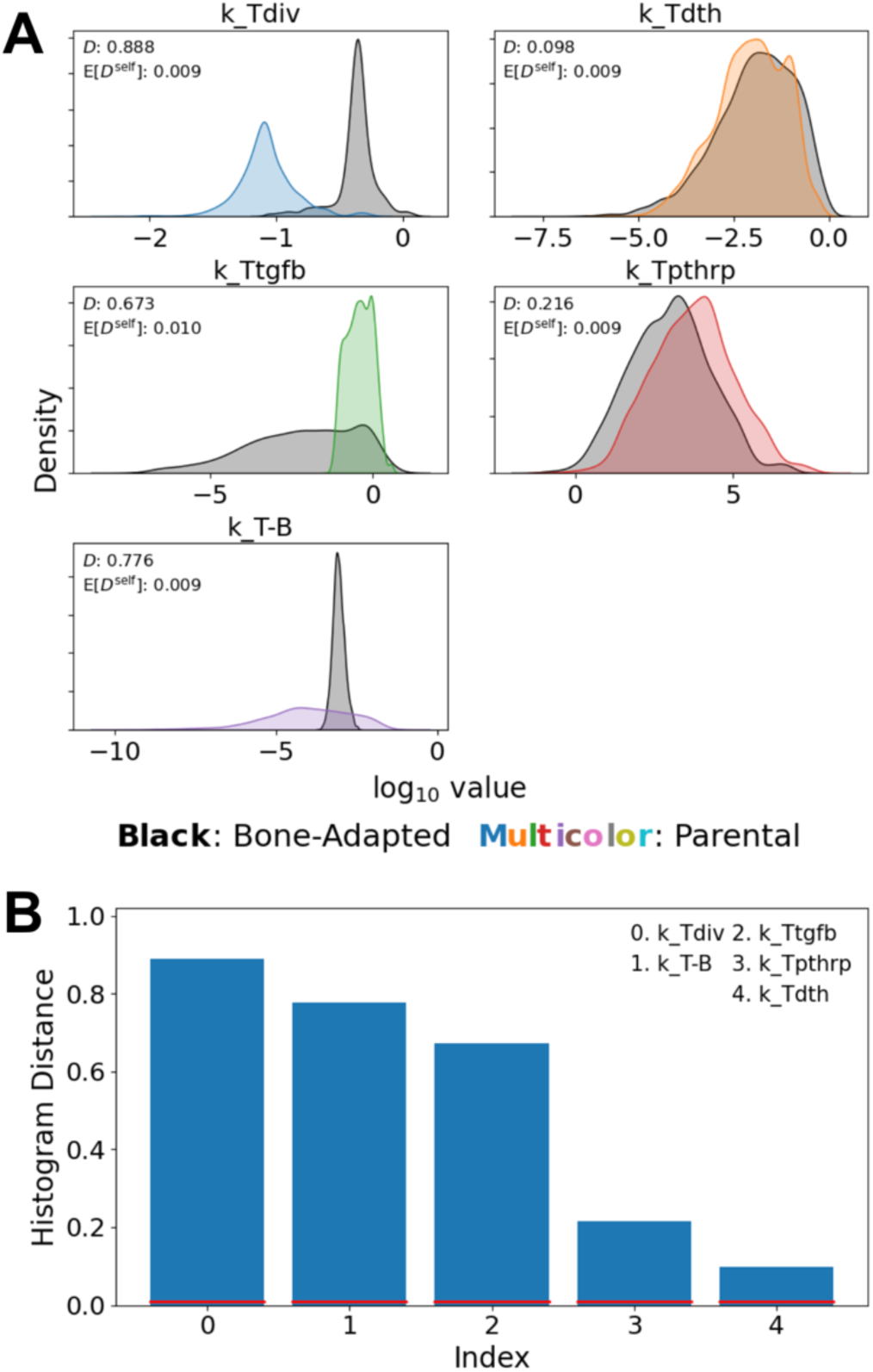
Comparisons of tumor-associated model parameters for the bone-adapted and non-adapted tumor types. **(A)** Distribution overlays for the five tumor-associated model parameters. Histogram distances, *D*, and self-distances, E[*D^self^*], are included to quantify the degree of overlap between distributions. **(B)** Parameters from *(A)* are rank ordered based on histogram distance. Red lines are values of the self-distance. In all cases, *D* ≫ E[*D^self^*], indicating statistically significant differences between distributions.

Parameter value differences between the fits to the bone-adapted and parental-derived tumor data are evident in both the means and widths of the histograms for several parameters. In particular, the largest difference is observed for the basal tumor division rate (*k_Tdiv_*), where the bone-adapted distribution is shifted toward higher values relative to the parental-derived distribution. For the TGF-β–enhanced rate of tumor cell division (*k_Ttgfb_*), the parental-derived distribution is narrowly concentrated over a limited range of values, illustrating the relative sensitivity of the non-adapted tumor to this parameter. Conversely, the bone-adapted distribution spans a broader range, demonstrating relative insensitivity to TGF-β. The rate of PTHrP production by tumor cells (*k_Tpthrp_*) exhibits substantial overlap between the bone-adapted and parental-derived tumor types, but with a clear shift in the mean toward higher values for the parental-derived tumor. This is an interesting result, since at first it may seem contradictory to the increased bone destructiveness of the bone-adapted tumor. However, since bone-adapted tumors grow significantly faster (Fig. 3, *bottom right*), they produce more *overall* PTHrP, even with lower per-cell rates of PTHrP production. The distributions for the tumor-induced OB death/differentiation rate parameter (*k_T–B_*) also showed pronounced differences, with a narrow and tightly constrained distribution for the bone-adapted tumor and a substantially broader distribution for the parental-derived tumor. Rather than reflecting a biological effect, however, this difference is likely due to increased uncertainty in the value of this parameter for the non-adapted tumor due to a lack of quantitative data for OBs in the Johnson et al. study (16) (Fig. 3, *bottom left*). Finally, the posterior distributions for the tumor death rate (*k_Tdth_*) largely overlap, indicating that differential dynamics between tumor types are not driven by differences in death rates.

### The Computational Model Predicts Differential OC and OB Dynamics Underlying Tumor-Type-Specific Bone Destruction

To test the predictive ability of the calibrated computational model, we used the inferred tumor-type-specific parameter ensemble to generate model predictions for OC and OB dynamics in the Johnson et al. (16) experimental setting, where direct measurements of bone-resident cell counts were unavailable (Fig. 5). Predicted OC dynamics for the parental-derived tumor show a monotonic increase in OC abundance over time, but at a substantially lower rate than that observed for the bone-adapted tumors used in this study (Fig. 5, *top right*). This difference is not unexpected and parallels the more gradual decline in bone density seen for the non-adapted tumor (Fig. 5, *top left*), indicating weaker overall OC-mediated bone resorption relative to the bone-adapted case. In contrast, the predicted OB dynamics for the Johnson et al. (16) experiment span a wide range of possible trajectories (Fig. 5, *bottom left*), including scenarios in which OB abundance decreases substantially, remains near baseline, and even slightly *increases* over time. This broad range of predicted outcomes reflects uncertainty in the parameter values due to the lack of experimental data on OB dynamics in the Johnson et al. dataset (16), underscoring the need for additional experimental measurements to resolve the underlying cellular dynamics.

**Figure 5.**
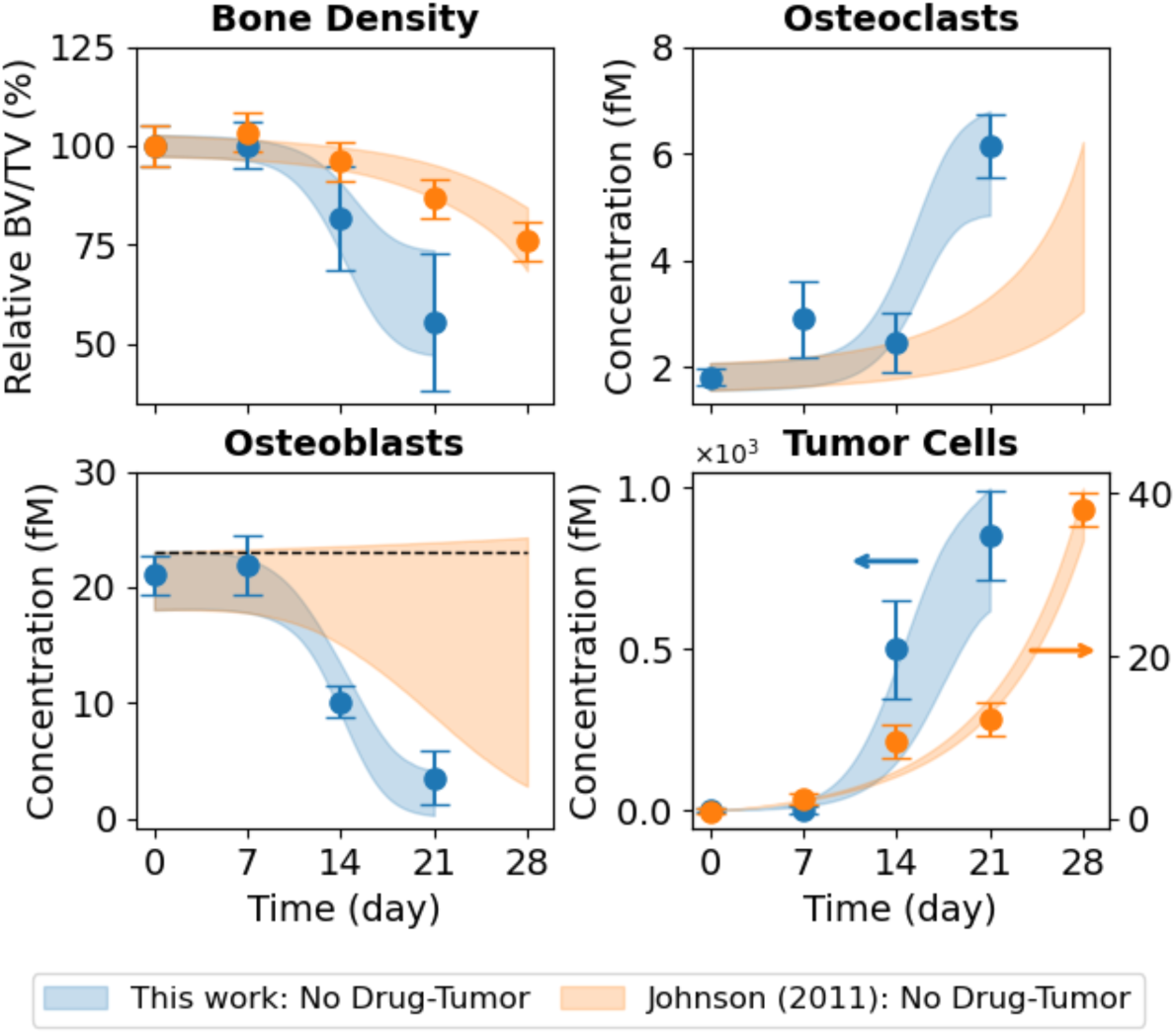
Predictions of OC and OB dynamics for the non-adapted vs. bone-adapted tumor type. *In vivo* data for the bone-adapted (this work; *blue*) and parental-derived non-adapted (Johnson et al. (16); *orange*) tumor types show bone density (*top left*) decreases in response to tumor growth (*bottom right*). However, bone destruction is more severe for the bone-adapted tumor. Using the inferred parameter sets (Figs. 4A and S3), the model predicts an increase in OCs for the non-adapted tumor (*top right*), but at a reduced rate relative to the bone-adapted tumor. For OBs (*bottom left*), due to parameter uncertainty, the model predicts a wide range of possible responses, from a slight increase to a significant decrease, albeit at a slower rate than the bone-adapted tumor. The black dashed line indicates zero change. In all plots, experimental error bars are standard errors (σ/√n) and model fits and predictions are shown as shaded 90% confidence envelopes. Bone density is plotted as a percentage relative to Day 0. BV/TV: bone volume ÷ tumor volume.

### Simulated OC Inhibition Stabilizes Bone but Yields Variable Effects on Bone-Adapted Tumor Growth

As an additional test of the predictive capability of our computational model, we used the calibrated parameter ensemble to simulate the effects of ZA treatment on bone remodeling and tumor dynamics in the bone-adapted setting (Fig. 6). Emulating the ZA-treatment experiments reported in Johnson et al. (16) (Fig. 3, *top left* and *bottom right*), ZA was introduced in the *in silico* experiments six days after tumor cell injection. Model simulations show that ZA treatment inhibits bone destruction, with predicted bone density stabilizing ∼100–130% of the initial value by 21 days post-treatment (Fig. 6, *top left*). This stabilization coincides with an expected rapid and substantial reduction in OC abundance following treatment (9) (Fig. 6, *top right*). Predicted OB dynamics showed little to no change relative to the untreated condition (Fig. 6, *bottom left*), indicating that bone protection due to ZA treatment arises in the model primarily through suppression of resorption rather than direct effects on OB populations. In contrast to the robust and consistent effects on bone and OCs, predicted tumor responses to ZA treatment span a wide range of outcomes (Fig. 6, *bottom right*), including trajectories in which tumor growth is largely unaffected and trajectories exhibiting partial or substantial attenuation of tumor expansion. This variability is consistent with the broad distribution for TGF-β–enhanced tumor growth rate parameter (*k_Ttgfb_*) for the bone-adapted tumor (Fig. 4). Altogether, these results indicate that while ZA treatment is predicted to reliably stabilize bone density for bone-adapted tumor-bearing mice through OC suppression, its effects on tumor growth are more uncertain, consistent with studies showing mixed effects on tumor burden (13–15).

**Figure 6.**
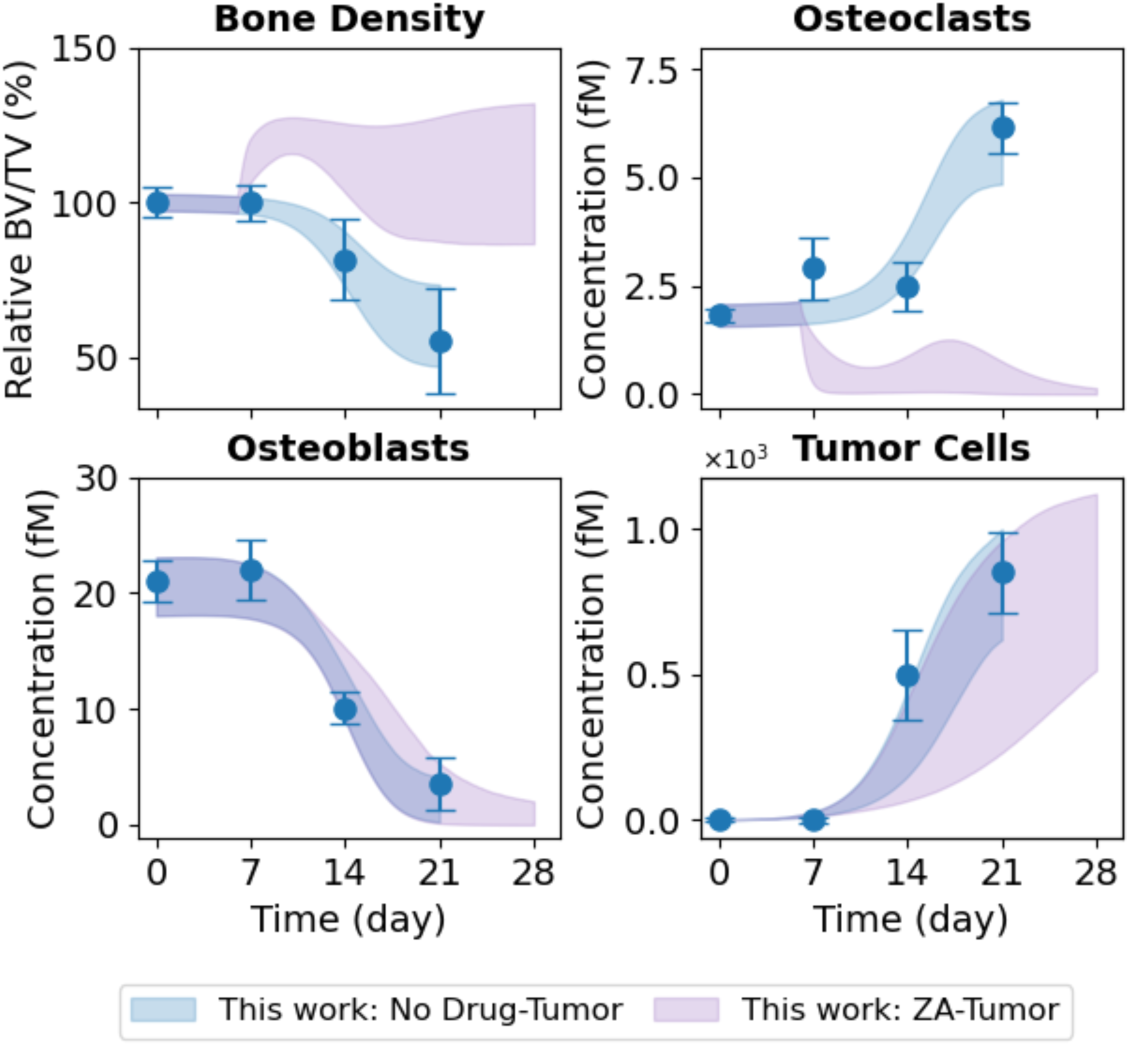
Predicted impact of drug treatment on bone density, OC, OB, and tumor cell dynamics for the bone-adapted tumor type. *In silico* experiments were performed with zoledronic acid (ZA) added six days after tumor cell injection. Using the inferred parameter sets (Figs. 4A and S3), the model predicts an initial spike in bone density (t*op left*), followed by a possible decline that levels out around Day 14. For OCs (*top right*), the model predicts a rapid decline followed by a possible temporary increase starting around Day 14, before eventually stabilizing near zero. No appreciable change in OB concentration is predicted (*bottom left*). For tumor cells (*bottom right*), the model predicts a range of possible drug responses, from no effect to a significant reduction in tumor growth. In all plots, experimental error bars are standard errors (σ/√n) and model fits and predictions are shown as shaded 90% confidence envelopes. Bone density is plotted as a percentage relative to Day 0. BV/TV: bone volume ÷ tumor volume.

## Discussion

In this study, we developed and calibrated a mechanistic population-dynamics model that integrates tumor cells, OBs, OCs, bone, and pharmacologic perturbation (Figs. 1, S1 and Tables 1, S1, S2) to investigate tumor-type-specific mechanisms underlying TIBD. By fitting the model to longitudinal *in vivo* data for tumors derived from both non-adapted parental and bone-adapted MDA-MB-231 breast cancer cells (Figs. 2, S2–S4), we obtained parameter sets that reproduce experimental trajectories of bone density, tumor burden, and bone-resident cell populations (Figs. 3, S5, S6). Comparative analysis of inferred tumor-associated parameter distributions revealed clear distinctions between tumor types (Fig. 4), most notably a higher basal tumor division rate and reduced sensitivity to TGF-β–enhanced proliferation in bone-adapted tumors, and greater TGF-β dependence and increased PTHrP production in parental-derived tumors. Model predictions further suggested that bone destruction associated with parental-derived tumors arises from a slower but meaningful increase in OC activity relative to bone-adapted tumors (Fig. 5). In contrast, due to limited experimental data, OB responses vary widely, from significant reduction to little or no change (Fig. 5). Simulated ZA treatment of bone-adapted tumors robustly stabilized bone density and suppressed OCs but yielded highly variable tumor growth dynamics (Fig. 6), elucidating the need for further experimental study of the effects of OC inhibition on bone-adapted tumor proliferation.

Together, the results presented here provide quantitative support for the central hypothesis of this work that tumor adaptation to the bone microenvironment reduces dependence on bone-derived growth factors and the efficacy of OC-targeted therapy. The strongest evidence comes from the comparative analysis of tumor-associated parameters (Fig. 4), which reveals systematic differences between non-adapted and bone-adapted tumors, particularly the larger basal division rate for bone-adapted tumors and the greater dependence on TGF-β–enhanced proliferation and increased PTHrP production for parental-derived tumors. As mentioned above, the latter result is notable because it seemingly contradicts the increased bone destructiveness of bone-adapted tumors (36), since PTHrP secretion is how tumor cells induce osteolysis. However, our analysis attributes the elevated bone destruction of bone-adapted tumors to the significantly higher proliferation rate relative to non-adapted tumors. This allows bone-adapted tumors to downregulate their *per-cell* PTHrP production while still maintaining high overall PTHrP levels due to higher cell counts. A rationale for this may be that it allows bone-adapted tumors to redirect energy towards alternative survival mechanisms, explaining their observed insensitivity to OC inhibition therapies (8, 10–12). Conversely, non-adapted tumors remain more dependent on bone-derived growth factors to compensate for their slower basal proliferation rate, resulting in increased sensitivity to disruption of the OC-driven vicious cycle. Importantly, note that these differences in parameter values between non-adapted and bone-adapted tumors were not imposed *a priori*, but emerged directly from the parameter inference process. Thus, our model recovered biologically feasible features of tumor adaptation to the bone microenvironment directly from experimental data.

The ultimate test of our central hypothesis is the impact of ZA treatment on tumor proliferation in the bone-adapted setting, where little to no effect is expected. However, our simulations show a broad range of predicted responses, from negligible impact to substantial tumor reduction (Fig. 6). Thus, our results do not provide direct support for our central hypothesis but, critically, also do not refute it. From a systems biology perspective, this result should be interpreted as revealing the limitations of mechanistic modeling in the face of limited data. On one hand, our approach has extracted maximum insight on factors underlying TIBD from the available *in vivo* data. At the same time, it has illustrated that the current data is insufficient for making precise predictions in various scenarios, such as OB dynamics during early stages of tumor adaptation (Fig. 5) and the effectiveness of OC-targeted therapies on bone-adapted tumors (Fig. 6). As such, our computational analysis can be viewed as advancing mechanistic understanding of TIBD while also exposing key knowledge gaps, which can be used to guide future experimental efforts under different experimental conditions. Toward this end, we note that the *in vivo* data generated here and in Johnson et al. (16) reflect limitations in experimental methods that can be addressed in future work. Specifically, the bone-adapted model used here is aggressive, which limits the data that can be obtained due to limited tissue availability in mice with aggressive tumors and exclusion of mice without successful tumor establishment (*Materials and Methods*). Similarly, as described above, the error bars from the Johnson et al. (16) data had to be modified for the sake of parameter identifiability (Fig. S4; *Materials and Methods*). Future applications of the computational model will be significantly strengthened by increasing the amount and reducing the uncertainty of the experimental data used for calibration.

Beyond biological insights, an additional contribution of this work is demonstrating the power of predictive, mechanistic modeling for studying complex diseases such as TIBD (37). The mechanistic model developed here is not simply a static representation of the bone microenvironment but serves as an extensible *in silico* testbed for hypothesis generation, experimental design, and therapeutic exploration. Future directions will add mechanistic detail to the model (both at the cell-population and intracellular levels), incorporate additional experimental data, and explore alternative therapeutic modalities and treatment schedules. This will be facilitated by the open and modular structure of the model, which enables direct reuse and extension by the broader community. We envision this framework as the foundation upon which an increasingly detailed and predictive model, or “Digital Twin” (38, 39), of the tumor–bone microenvironment will be built, leading to tighter integration between experimental and computational efforts and accelerating discovery in the study of TIBD.

## Materials and Methods

### Cell Culture

Human triple-negative breast cancer cells (MDA-MB-231) were obtained from the American Type Culture Collection. A bone-adapted derivative (MDA-MB-231b) was established through *in vivo* selection for enhanced bone tropism, as previously described (7, 11, 14, 25, 40). Cells were maintained in Dulbecco’s Modified Eagle Medium (high glucose, pyruvate; Gibco) supplemented with 10% fetal bovine serum (Peak Serum) and 1% penicillin/streptomycin (Corning) under standard humidified culture conditions (37 °C, 5% CO_2_). Cultures were routinely tested for mycoplasma contamination and used at low passage to preserve the bone-adapted phenotype.

### Murine Model of Bone Metastasis

Bone metastasis was modeled using female athymic Nude-Foxn1^nu^ mice (7.5 weeks old; Envigo/Inotiv), an immunocompromised strain recommended for human breast cancer cell engraftment. A total of 48 mice were used, half in the control group and half injected intratibially with tumor cells. Both groups were further subdivided into three cohorts of eight, with scheduled euthanasia at days 7, 14, and 21 post-injection. Mice in the tumor cohort were anesthetized with continuous isoflurane inhalation and injected with 50,000 MDA-MB-231b cells suspended in 10 µL sterile phosphate-buffered saline (PBS; Gibco) using a 29-gauge needle, as previously described (11, 12, 14, 40). Control mice received PBS alone. Animals were monitored biweekly for weight and general health. Disease progression was evaluated for each mouse by radiography immediately prior to sacrifice. Right tibias were collected for imaging and histological analysis.

### Radiographic Imaging and Lesion Quantification

Radiographs were acquired using a Faxitron LX-60 digital X-ray system at 35 kVp for 8 s. Mice were anesthetized under continuous isoflurane inhalation and laid in the prone position. Osteolytic areas were manually segmented in ImageJ (41). Animals without measurable lesions at later timepoints (14 and 21 days) were excluded from analyses. All quantitative plots and statistical tests were generated using GraphPad Prism 10.2.0 (GraphPad Software, Boston, MA USA).

### Micro-Computed Tomography (µCT)

At sacrifice, tibias were fixed in 10% neutral buffered formalin (Sigma) for 48 hours at 4°C. Following fixation, tibias were transferred to 70% ethanol and scanned using a Scanco µCT40 system (70 kVp, 12 µm voxel size, 300 ms integration time). These acquisition parameters are consistent with recommended settings for evaluating trabecular degradation and cortical destruction in bone metastasis models. Images were reconstructed with Gaussian filtering and thresholded using consistent global settings across samples (Gauss σ = 0.2, Gauss support = 1.0, lower threshold = 190, upper threshold = 1000). Bone morphometric parameters were quantified within a standardized region located 30 slices below the proximal growth plate and extending through the next 100 slices. Bone morphometric parameters of the analyzed segments were calculated using the Scanco software.

### Histology and Cell Quantification

After µCT imaging, tibias were stored at 4°C in 70% ethanol and sent to the Vanderbilt University Medical Center (VUMC) Translational Pathology Shared Resource (TPSR) for tissue processing and H&E staining. Tibias were decalcified using Immunocal formic acid decalcifier (StatLab) overnight. Specimens were embedded in paraffin wax, sectioned at 5 µm, sliced on a microtome, stained with H&E, and coverslipped. Two H&E slides per mouse were examined under a microscope (Keyence BZ-X810) and imaged immediately distal to the growth plate at 10X magnification. Bone perimeter was manually outlined using the freehand line tool in ImageJ (41). OBs were identified as cuboidal basophilic cells lining mineralized surfaces with at least three cells in a row. Tumor boundaries were manually outlined, and cell counts were extracted using color-thresholding in ImageJ. Unstained mounted slides from TPSR were then stained for TRAP utilizing a substrate incubation step with Naphthol AS-BI (Sigma) in acetate buffer with tartaric acid (Sigma), followed by a color reaction using pararosaniline dye (Sigma) and sodium nitrite (Fisher). Two sections per mouse were counterstained with hematoxylin, coverslipped, and imaged as described above. OCs were defined as TRAP-positive multinucleated cells containing three or more nuclei. All slides were imaged using a Keyence digital microscope.

### In Vivo Experimental Data Normalization

Cell counts for tumor cells, OBs, and OCs were collected from histological staining (Fig. 2C,D) for up to two sections per mouse to capture variability due to location of each histological section within the bone. Due to aggressive disease in some mice in the tumor cohort, not every mouse could be represented in all counts due to limited slide availability. All cell counts were tabulated from at least seven slides for no less than four mice per group. To connect these data to the computational model developed here (Fig. 1), cell counts are converted to concentrations (pM) using the following relation,

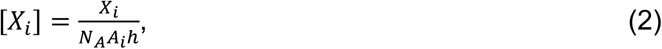

where [*X_i_*] is the cell concentration (tumor cells, OBs, or OCs) from slide *i*, *X_i_* is the raw cell count, *N_A_* is Avogadro’s number, *A*_i_ is the area of the “region of interest” for slide *i*, and *h* is the slide thickness (5 µm). For OBs and OCs, further normalization is necessary to account for the bone perimeter in each slide, which is assumed to be a random variable (i.e., slides with more bone perimeter, by chance, tend to have higher cell counts). Thus, we scale the OB and OC concentrations for each slide by the ratio of the average bone perimeter for all slides to the bone perimeter for each individual slide, i.e.,

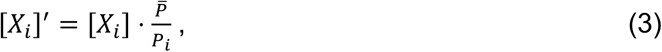

where [*X_i_*]’ is the normalized OB or OC cell concentration, *P̅* is the average bone perimeter, and *Pi* is the bone perimeter for slide *i*.

Bone volumes obtained from µCT (one measurement per mouse) are also normalized for use in the computational model. The bone volume/tumor volume (BV/TV) output for each mouse is divided by the average BV/TV for mice with no tumor by histology. These include all non-tumor-bearing mice (control cohort) and mice from the Day 7 tumor cohort (no tumor cells detected in all cases). This normalization gives an initial relative bone density of 100% (see Figs. 3, 5, and 6, *top left*). All *in vivo* data (raw and processed) are provided as Supporting Information (Dataset S1).

### Computational Model Construction and Simulation

The mechanistic model of TIBD presented in this work (Fig. 1) was constructed using PySB (42), an open-source Python-based platform for building, simulating, and analyzing models of complex biological systems. PySB is based on the rule-based modeling formalism (43, 44), in which models are specified in terms of “rules” that govern the basic processes (e.g., birth, death, differentiation) that occur within a biological system. Coupled sets of ordinary differential equations (ODEs) are automatically generated from these rules using BioNetGen (45, 46), an open-source Perl/C++ platform accessible through PySB. These ODEs can then be solved numerically in PySB using solvers available in the Python library SciPy (47).

The computational model includes *six species*: responding OBs (*R*), active OBs (*B*), active OCs (*C*), bone (*N*), tumor cells (*T*), and the bisphosphonate ZA (*Z*). These species participate in *12 reactions* (Table S1) with rate laws involving *four algebraic functions* (Table S2) and *36 constant parameters* (Table 1). Three of the species, five reactions, three functions, and 23 parameters come directly from Lemaire et al. (5). The additional model elements were added in this work to model bone, tumor, and ZA dynamics and impacts on bone-resident cell populations. Responding and active OBs, active OCs, and tumor cells are represented in the model in concentration units (fM). Bone, on the other hand, is represented in generic “bone units”, such that the reaction *B* → *B* + *N* (Table S1, reaction 6) should be read as “an active OB deposits one unit of bone.” The initial amount of bone can be set to any value depending on how much bone one wants to assume each active OB deposits (or OC resorbs; Table S1, reaction 7). Bone density is then compared to experimental values (in relative percent) by dividing the amount of bone at any time point by the initial value and multiplying by 100. Finally, ZA is represented within the model as a ratio with respect to the amount (0.1 mg/mouse) used in the experiments reported by Johnson et al. (16). The rationale for this is we do not have any information about drug bioavailability (i.e., how much drug actually reaches the tumor) or the volume of the tumor microenvironment. This is a reasonable and common approach in systems biology/pharmacology modeling that simply requires the modeler to recognize that a value of 1 corresponds to the reference drug concentration.

Model simulations were performed using VODE (48), a well-established numerical ODE solver accessible in PySB through the Python package SciPy (47). In all cases, the initial concentrations of responding and active OBs, active OCs, and tumor cells were set to zero. Initial amounts of bone and ZA were set to 100 and zero, respectively. Pre-equilibration simulations were run for 500 simulated days to allow the system to reach steady-state levels of OBs, OCs, and bone density for a given parameter set. *In silico* experiments were then performed emulating the *in vivo* experiments reported here (Fig. 2) and in Johnson et al. (16). Specifically, three types of experiments were simulated: (i) tumor cells injected at Day 0 (untreated, tumor-bearing mice); (ii) tumor cells injected at Day 0 followed by injection of ZA at Day 6 (ZA-treated, tumor-bearing mice); and (iii) ZA injected at Day 6 (ZA-treated, non-tumor-bearing mice). The amount of injected ZA was set to 1 to match that used in the Johnson et al. (16) experiments (i.e., 0.1 mg/mouse). The concentration of injected tumor cells (1 fM) was chosen because it is smaller than the lowest measured tumor cell concentration in our experiments (∼5.5 fM; see Dataset S1), which occurred at Day 14. No tumor cells were detected in any mice prior to Day 14, even though we know tumor cells were present, since they eventually proliferated to detectable levels. Thus, setting the injected concentration to be lower than the lowest detected level is a reasonable assumption. Note this assumption can be relaxed in future studies by allowing the injected tumor concentration to be a free parameter estimated during model calibration.

### Monte Carlo-Based Parameter Estimation

PySB is closely associated with PyDREAM (32), an open-source Python implementation of the DiffeRential Evolution Adaptive Metropolis (DREAM) algorithm (49, 50), a Markov Chain Monte Carlo (MCMC) method for parameter estimation. PyDREAM is a multi-chain algorithm where multiple MCMC walkers are instantiated in parallel, with intermittent communication between chains (49, 50). Parameter convergence is quantified using the Gelman-Rubin (GR) metric (33), a measure of the ratio of the variability of the sampled parameter values across the chains to that within the chains. Convergence is achieved when GR values for all parameters are close to 1.

Here, we used PyDREAM with five MCMC chains to sample 30 of the 36 model parameters (see Table 1). All 30 parameters were sampled in log-space with Bayesian priors defined as log-normal distributions centered on the initial parameter values (Table 1), except for the carrying capacity *M*, which was assigned a log-uniform prior centered on its initial value with a range of 0.3. We set the GR convergence threshold to 1.2 and initialized PyDREAM to run for 50,000 iterations. At each iteration, the *in silico* experiments described above were run to compare to the *in vivo* data generated here (Fig. 2) and extracted from Johnson et al. (16). The quality of the fit was quantified as the sum of the log-likelihoods over all data points (Fig. S5), with larger values corresponding to better fits (32). After 50,000 iterations, PyDREAM performed a convergence check and if convergence was not achieved (i.e., any GR value > 1.2), the run was automatically continued for an additional 50,000 iterations. This process was repeated until convergence was reached. For the results presented here, PyDREAM ran for 500,000 iterations before reaching convergence (Fig. S5), generating a total of 2.5 million parameter samples. The first half of the iterations were discarded, considered as “burn-in,” leaving 1.25 million samples. The parameter set ensemble was further filtered by calculating the mean value of the log-likelihoods over these samples and discarding any sets with log-likelihood values below the mean value less two standard deviations (Fig. S5). All PyDREAM runs were executed in parallel on 32 processors on two dedicated Dell PowerEdge workstations, equipped with dual AMD EPYC 7543 processors (64 cores) and 1 TB memory, housed and maintained at the Arkansas High-Performance Computing Center (AHPCC).

### Extraction and Processing of Bone Density and Tumor Burden Data for Parental-Derived Tumors from Published Figures

To aid model calibration, specifically estimation of the rate parameter for ZA-mediated OC inhibition (*k_Z–C_*; Tables 1 and S1), bone volume and tumor burden data for *in vivo* experiments using parental MDA-MB-231 cells were extracted from Figures 4D and 5D of Johnson et al. (16). This was done using the WebPlotDigitizer (version 5) (51, 52), an online tool that uses machine learning methods to extract quantitative data from published visualizations. For bone volume, data is reported at Days 7, 14, 21, and 28 post-injection for four separate conditions: (i) untreated, non-tumor-bearing, (ii) untreated, tumor-bearing, (iii) ZA-treated, non-tumor-bearing, and (iv) ZA-treated, tumor-bearing. To convert these data to relative bone densities, we used the mean value of the untreated, non-tumor-bearing data as the baseline and divided the data points for the other three datasets by this value. This gave us three datasets, all initialized at 100% bone density, for calibrating the computational model (see Fig. 3, *top left*).

For tumor burden, Johnson et al. (16) report fluorescence data for untreated and ZA-treated tumor-bearing mice at Days 14, 21, and 28 post-injection. To aid parameter calibration, we estimated fluorescence values at Days 0 and 7 for the untreated data by fitting an exponential curve to the data points at Days 14, 21, and 28 (Fig. S3). We then fit an exponential curve to the ZA-treated data, forcing the pre-exponential factor to be the same as for the untreated data (Fig. S3), since this value represents the estimated fluorescence at Day 0 when tumor cells are injected (the same number of cells are injected in both cases). Importantly, we chose to exclude the ZA-treated data points at Days 14 and 21 in the fit, using only the data point at Day 28 because the fluorescence values for the other points are very similar, indicating they may be below the limit of detection (Fig. S3). Thus, fluorescence values for ZA-treated tumors at Days 0, 7, 14, and 21 were estimated from this exponential fit. Finally, since we assume 1 fM of tumor cells are injected at Day 0 in both cases (see above), tumor cell concentrations were estimated for subsequent days by dividing the measured or estimated fluorescence values by the Day 0 fluorescence.

### Modification of Error Bars from Published Data

Initial attempts to calibrate the computational model (Fig. 1) using the bone density and tumor burden data from Johnson et al. (16) were complicated by large error bars in the data. These error bars made it difficult for PyDREAM (32) to infer parameter sets that distinguish the bone density and tumor burden values at different time points. Therefore, to facilitate parameter estimation, we reduced the sizes of the Johnson et al. (16) error bars, while maintaining the mean values (Fig. S4). Specifically, for bone density, we set all error bars equal to 5 (relative percent); for tumor burden, error bars were set to 5% of the mean value for each data point (measured and estimated; see Fig. S3).

### Quantifying Differences Between Posterior Parameter Distributions

MCMC-based parameter estimation methods, such as PyDREAM (32), produce many parameter sets that closely match experimental data, also known as an “ensemble,” rather than a single “best-fit” parameter set (53). As such, parameter comparisons must account for the *distributions* of parameter values across the inferred ensemble. These marginal distributions, known as “posteriors” (53), can span many orders of magnitude and have arbitrary shapes (e.g., normal, long-tailed, bimodal). Thus, comparisons using point statistics, such as mean and standard deviation, can miss important aspects of the distributions being compared.

To compare posterior parameter distributions more comprehensively, we used the “histogram distance” (34, 35) (see Fig. 4), defined as

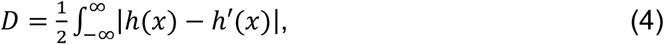

where *h*(*x*) and *h’*(*x*) are the values of the two distributions at *x*. Since both distributions are normalized (i.e., sum to 1), the factor of ½ ensures that *D* ranges between 0 and 1, with 1 indicating no overlap between distributions and 0 a perfect match. That the value of *D* conveys information about the extent of separation between distributions is what makes it particularly appealing relative to alternatives, like the Kolmogorov-Smirnov metric (34), which enable hypothesis testing but do not provide intuitive measures of difference. Additionally, we calculate the “self-distance,” a measure of intrinsic variability in the calculation of a distribution (34, 35). The self-distance is defined as in Eq. (4) but with *h*(*x*) corresponding to a distribution based on a finite number of samples and *h’*(*x*) the theoretical *true* distribution based on an infinite number of samples. The expected value of the self-distance can then be estimated as (35)

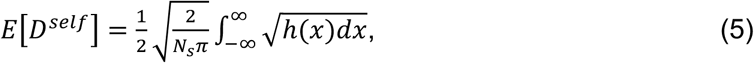

where *N_s_* is the number of parameter samples (note that *dx* is under the square-root). This value corresponds to the expected deviation of a distribution from ground truth and can thus be used as a statistical threshold for significance, i.e., if *D* < E[*D^self^*] there is no statistical evidence the two distributions are different (see Fig. 4B). As expressed in Eqs. (4) and (5), *D* and E[*D^self^*] are defined for continuous distributions. Here, we generate continuous posterior parameter distributions from PyDREAM samples using the ‘kdeplot’ function (with ‘bw_adjust=3’) in the ‘seaborn’ Python package (54). *D* and E[*D^self^*] are then calculated numerically by dividing the range of *x* values into a minimum of 1000 segments and estimating the integrals as discrete summations.

## Code Availability

All PySB model files, parameter calibration scripts (with experimental data), and analysis codes are freely available at github.com/SysBioCollab-UArk/TIBD_microenvironment. Parameter calibrations were run using the ‘pydream-rewrite’ branch of the PyDREAM source code, available at github.com/LoLab-MSM/PyDREAM/tree/pydream-rewrite.

## Ethics Statement

All animal protocols were approved by Vanderbilt University Institutional Animal Care and Use Committee and were conducted according to National Institutes of Health guidelines for care and use of laboratory animals.

## Figures

**Figure S1.**
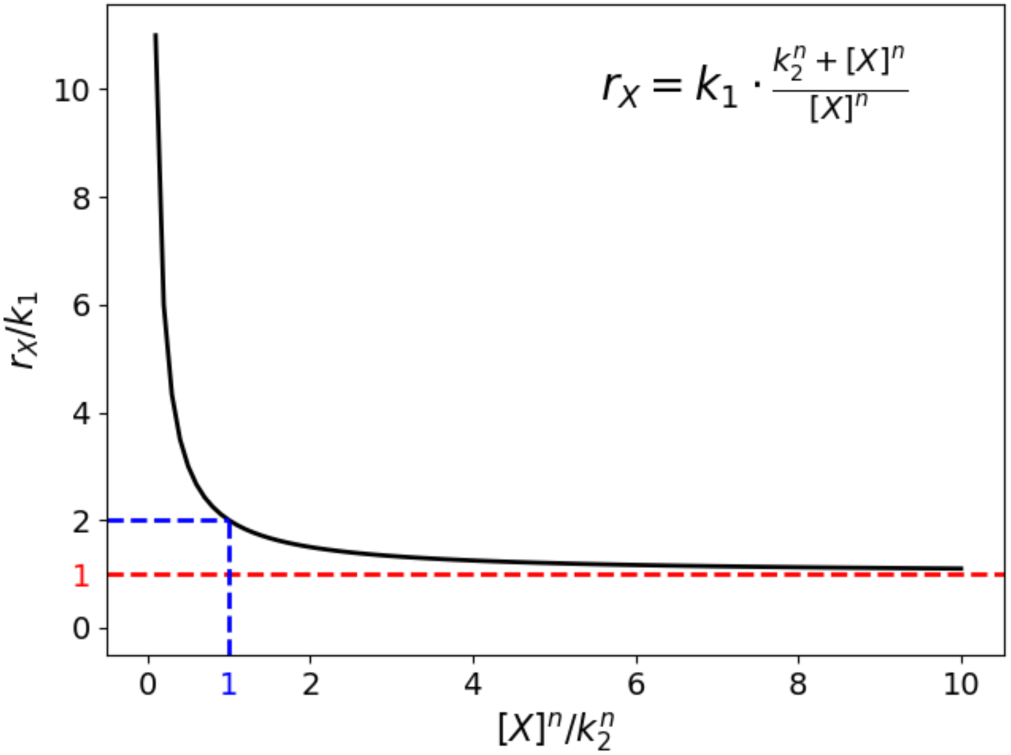
Illustration of the per-cell reaction rate expression used to model OB bone formation and OC bone consumption. The per-cell reaction rate, *r_X_*, is shown in the top-right corner of the plot, while the axis values are scaled to non-dimensionalize the equation. The red dashed line indicates that at large values of [*X*] (OBs or OCs), *r_X_* → *k_1_*; the blue dashed line indicates that *r_X_* = 2*k_1_* when [*X*] = *k_2_*. Square brackets represent concentration.

**Figure S2.**
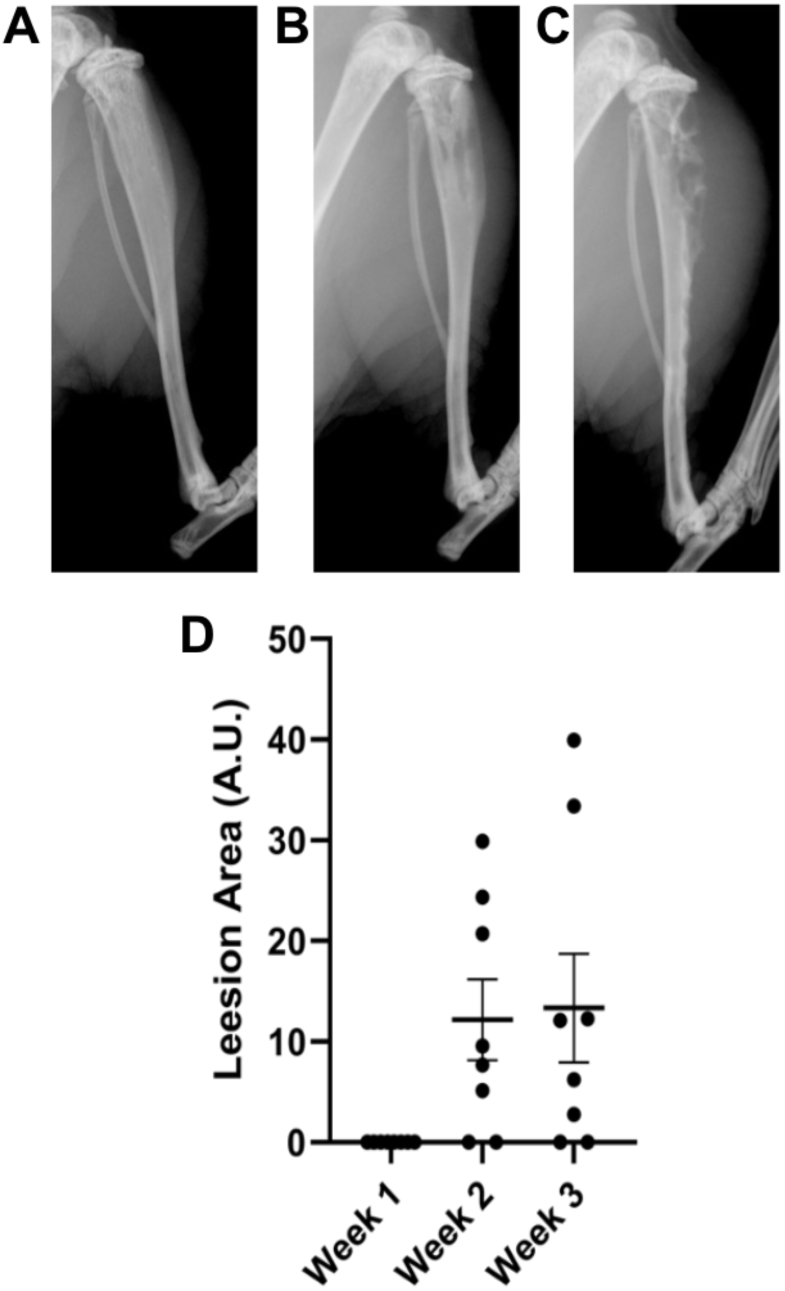
Representative x-ray images of mice after tumor injection. X-ray images show tibias of mice one week **(A)**, two weeks **(B)**, and three weeks **(C)** after MDA-MB-231b (bone-adapted) injection. **(D)** Quantification of lesion area (mean ± standard error). Mice with no lesions at two and three weeks after tumor cell injection were excluded from all downstream analyses. n = 8 per group.

**Figure S3.**
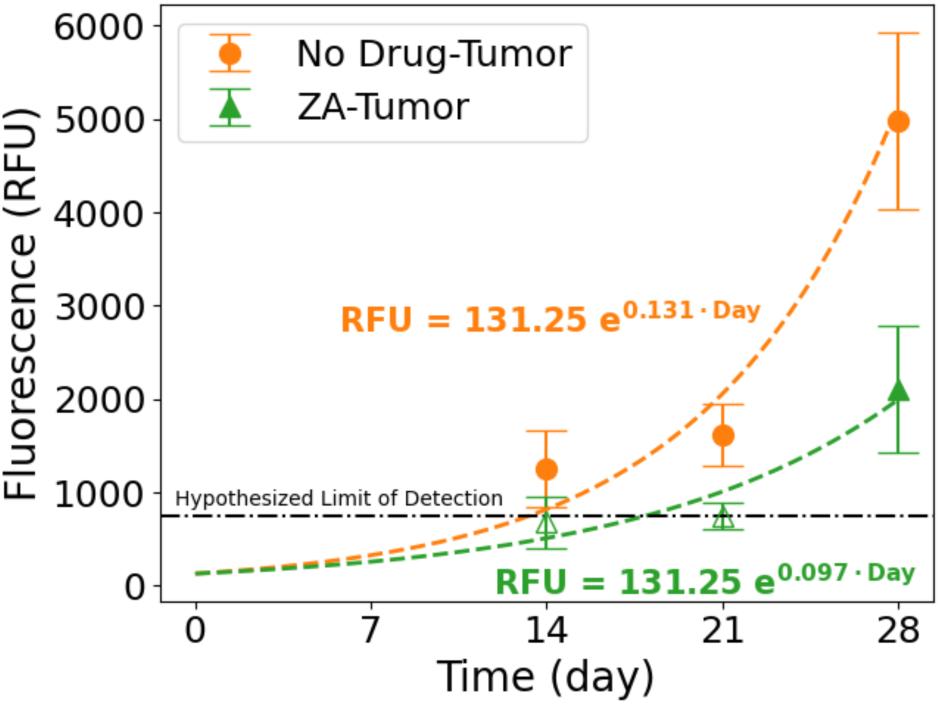
Exponential fits to fluorescence tumor burden data from Johnson et al. (16). Exponential prefactors for both datasets are purposely set equal since the number of injected tumor cells at Day 0 is the same in both cases. Also, data points for the ZA-treated tumor at 14 and 21 days post-injection (unfilled triangles) are excluded from the exponential fit since they fall below our hypothesized limit of detection (750 RFU). Assuming the injected tumor concentration is 1 fM at Day 0, these curves are used to estimate tumor cell counts at Day 7 for the untreated tumor (“No Drug–Tumor”) and Days 7, 14, and 21 for the ZA-treated tumor (“ZA-Tumor”; *Materials and Methods*). RFU: relative fluorescence units; ZA: zoledronic acid.

**Figure S4.**
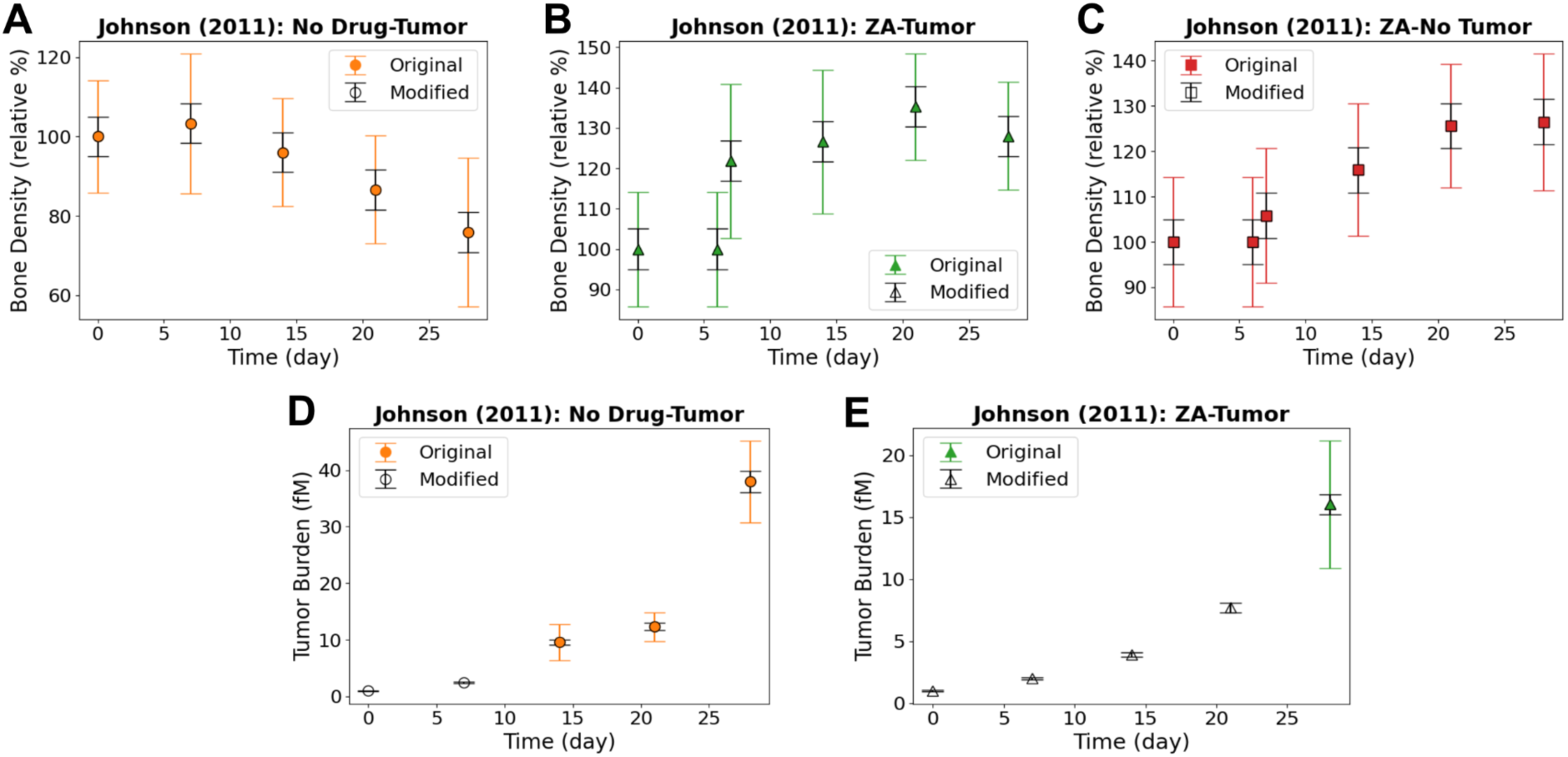
Comparison of original and modified error bars for the extracted datasets from Johnson et al. (16). **(A–C)** Bone density for untreated, tumor-bearing (*A*), drug-treated, tumor-bearing (*B*), and drug-treated, non-tumor-bearing (*C*) mice; **(D, E)** tumor burden for untreated, tumor-bearing (*D*) and drug-treated tumor-bearing (*E*) mice. Note that the “Modified” data points at Days 0 and 7 in panel *D* and Days 0, 7, 14, and 21 in panel *E* are estimated based on exponential fits to the original data points (see Fig. S3 and *Materials and Methods*). Thus, there are no “Original” data points for these cases.

**Figure S5.**
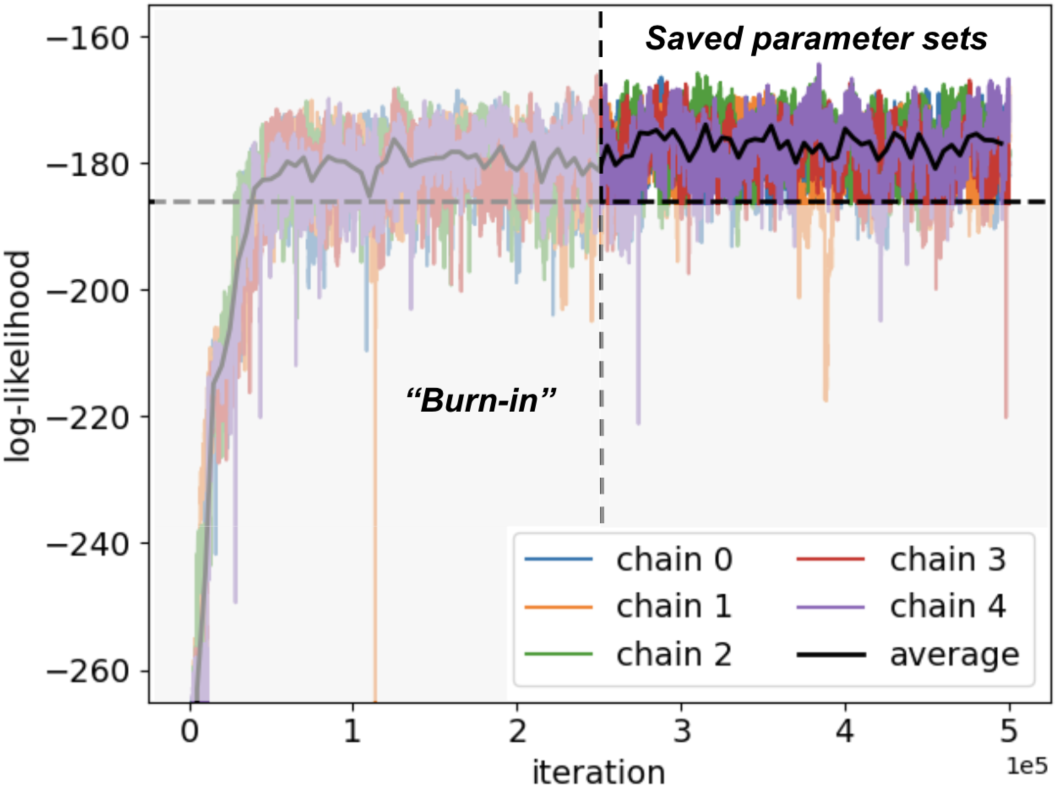
Log-likelihoods at each iteration for all five Markov-chain Monte Carlo (MCMC) chains during the PyDREAM calibration run. The run took a total of 5×10^5^ iterations. The first 2.5×10^5^ iterations were considered “burn-in” and discarded. The dashed horizontal line represents two standard deviations from the mean value of the log-likelihood, calculated over the last 2.5×10^5^ iterations. Parameter sets with log-likelihood values below this line were discarded. In total, 1,212,105 parameter sets, of which 81,467 are unique, were retained after filtering.

**Figure S6.**
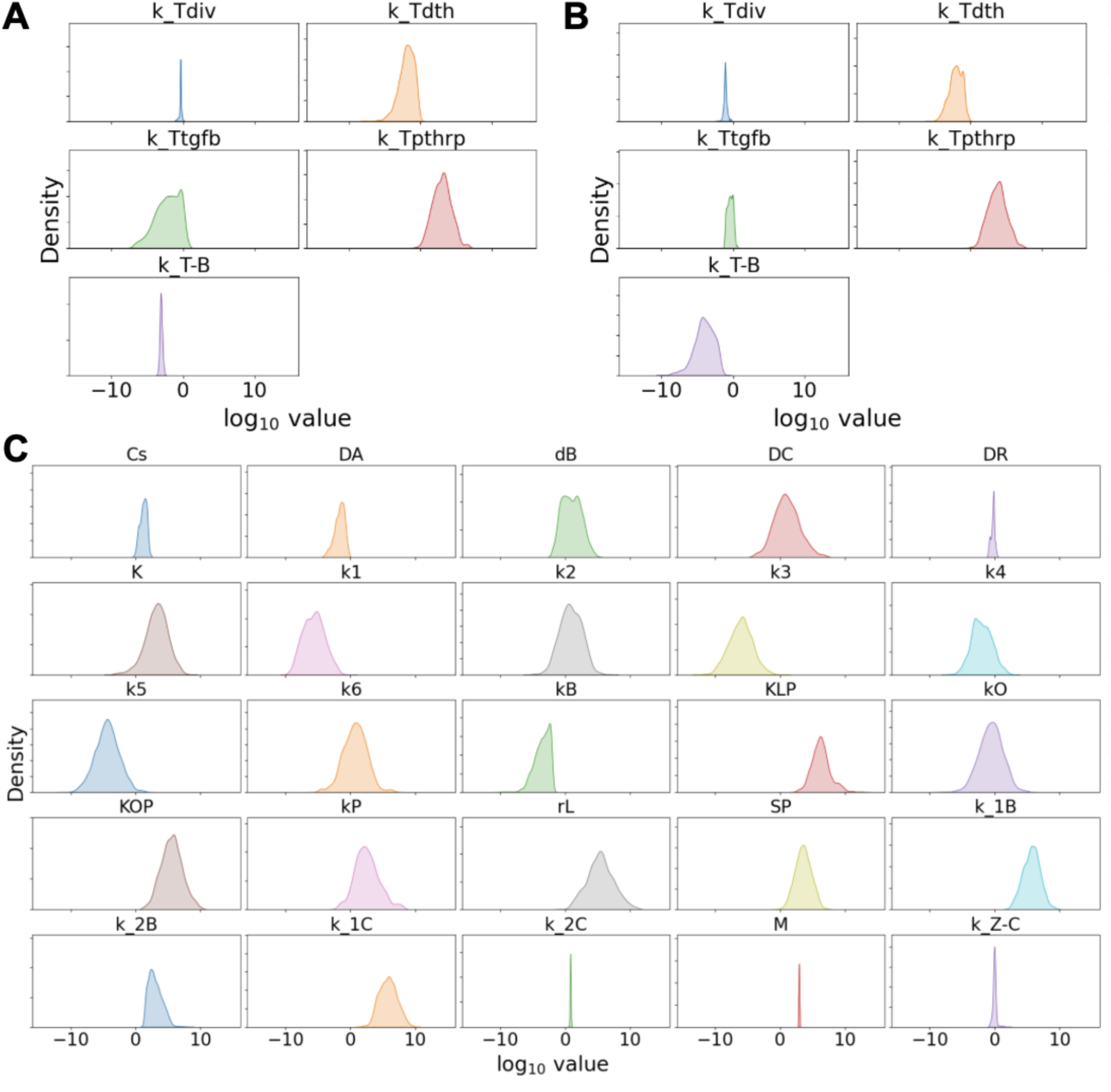
Distributions of parameter values inferred by PyDREAM. **(A, B)** The five tumor-associated model parameters for the *(A)* bone-adapted tumor (*this work*; see Figures 3, 5, and 6 of the main text) and *(B)* parental-derived non-adapted tumor (16) (see Figures 3 and 5 of the main text). Note that the distributions in *A* and *B* are overlaid in Figure 4A of the main text. **(C)** The 25 non-tumor-associated model parameters. All distributions are based on the 1,212,105 parameter sets sampled by PyDREAM. The x-axes are shared among all plots in *A–C*.

## Tables

**Table S1.**
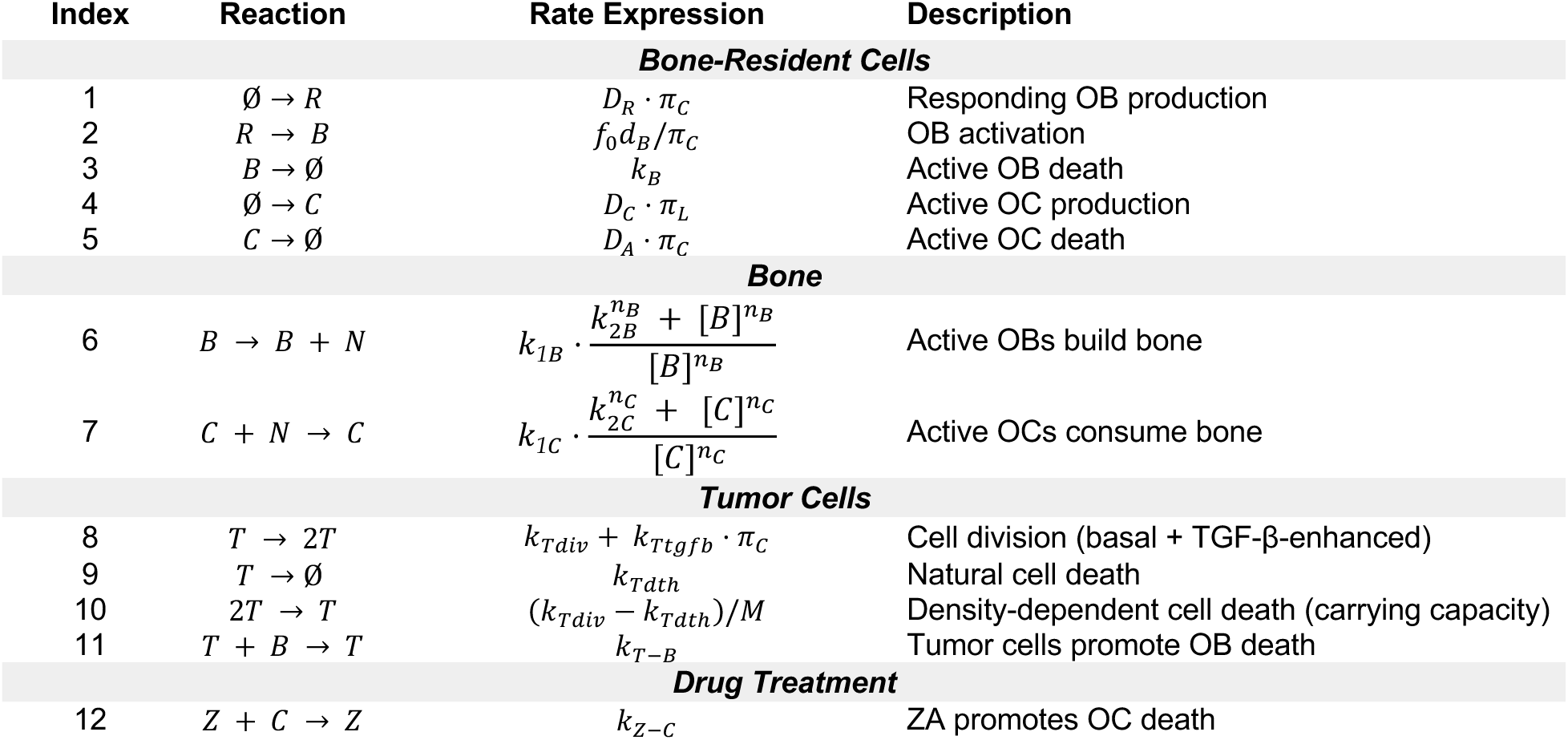
Mechanistic model of OB, OC, tumor, and bone dynamics. Reactions 1-5 are from the Lemaire et al. bone homeostasis model (5). Reactions 6-12 were added in this work to capture interactions between tumor, bone, osteoblasts (OBs), and osteoclasts (OCs), as well as the effect of drug treatment with zoledronic acid (ZA). *R*: responding OB, *B*: active OB, *C*: active OC; *N*: bone density; *T*: tumor cell; *Z*: ZA. Square brackets indicate concentration.

**Table S2.** Functions used in the rate expressions in Table S1. Functions 1-3 are from the Lemaire et al. bone homeostasis model (5). Function 4 was added in this work to capture PTHrP secretion from tumor cells. *R*: responding osteoblast (OB), *B*: active OB, *C*: active osteoclast (OC); *T*: tumor cell. Square brackets indicate concentration.

| Index | Function | Units | Description |
| --- | --- | --- | --- |
| **Bone-Resident Cells** |  |  |  |
| 1 | $\pi_P = \frac{I_p/k_p + S_p/k_p}{I_p/k_p + k_6/k_5}$ | unitless | Fraction of bound PTH receptors on ROB and AOBs |
| 2 | $\pi_C = \frac{f_0 C^s + [C]}{C^s + [C]}$ | unitless | Fraction of bound TGF-β receptors on uncommitted OB progenitors |
| 3 | $\pi_L = \frac{k_3}{k_4} \cdot \frac{K_L^P \cdot \pi_P \cdot [B]}{1 + \frac{k_3 K}{k_4} + \frac{k_1}{k_2 k_O} \cdot \left( \frac{K_O^P}{\pi_P} [R] + \bar{I}_O \right)} \cdot \left( 1 + \frac{\bar{I}_L}{r_L} \right)$ | unitless | Ratio of bound to unbound RANK receptors on OC precursors |
| **Tumor Cells** |  |  |  |
| 4 | $I_P = \bar{I}_P + k_{Tpthrp} \cdot \pi_C \cdot [T]$ | pM·day<sup>-1</sup> | Rate of injected PTH/PTHrP, external and from tumor |

## Data Availability

All data generated and analyzed in this study are included in the Supplementary Information (Dataset S1) and at github.com/SysBioCollab-UArk/TIBD_microenvironment.

## Author Contributions

A.G.V.: Conceptualization, formal analysis, investigation, visualization, writing–original draft, writing–review and editing.

N.E.B.: Conceptualization, data curation, investigation, methodology, validation, visualization, writing–original draft, writing–review and editing.

E.P.B.: Data curation, investigation, methodology, writing–review and editing. S.A.: Conceptualization, investigation, writing–review and editing.

R.C.: Formal analysis, investigation, writing–original draft.

A.N.: Formal analysis, investigation, writing–original draft.

H.V.: Formal analysis, investigation, writing–original draft.

A.J.: Formal analysis, investigation, writing–original draft.

J.A.R.: Conceptualization, funding acquisition, methodology, project administration, supervision, writing–review and editing.

L.A.H.: Conceptualization, formal analysis, funding acquisition, investigation, methodology, project administration, software, supervision, validation, visualization, writing–original draft, writing–review and editing.

## Acknowledgements

This work was funded by the National Institutes of Health (NIH) under awards K22CA237857 (L.A.H.), T32GM007347 (N.E.B.), and T32GM152284 (N.E.B.). Additional funding (to L.A.H.) was provided by the Arkansas Biosciences Institute and the University of Arkansas Women’s Giving Circle. A.G.V. thanks the University of Arkansas Interdisciplinary Graduate Program in Cell & Molecular Biology (CEMB) for funding support. L.A.H. thanks J.C. Pino for technical help with PyDREAM and acknowledges the NIH-sponsored “Multiscale Modeling and Viral Pandemics” working group and organizers of the “Building Immune Digital Twins” workshop hosted at the Institut Pascal, Université Paris-Saclay from May 15–June 2, 2023. The VUMC TPSR is supported by NIH/NCI Cancer Center Support Grant P30CA068485. This research is supported by the Arkansas High Performance Computing Center (AHPCC), which is funded through multiple National Science Foundation grants and the Arkansas Economic Development Commission.

